# Rab11FIP2 controls NLRP3 inflammasome activation through Rab11b

**DOI:** 10.1101/2025.05.19.654879

**Authors:** Caroline Gravastrand, Maria Yurchenko, Stine Kristensen, Astrid Skjesol, Carmen Chen, Zunaira Iqbal, Karoline Ruud Dahlen, Unni Nonstad, Liv Ryan, Terje Espevik, Harald Husebye

## Abstract

Membrane trafficking through the trans-Golgi network has recently been shown to guide activation of the NLRP3 inflammasome. The GTPases Rab11a and Rab11b, and their effector molecule Rab11-FIP2, are regulators of endosome trafficking and retrograde transport. Rab11-FIP2 binds phosphatidylinositol species including PI4P, enriched in the *trans*-Golgi network and peripheral endosomes following NLRP3 inflammasome activation. We here demonstrate that Rab11-FIP2 and Rab11b, but not Rab11a, control caspase-1 mediated cleavage of pro-IL-1β and GSDMD, and pyroptotic cell death in human macrophages. Rab11-FIP2 also controlled LPS stimulated IKKβ activation by TAK1 and IKKβ mediated NLRP3 translocation to the *trans*-Golgi network. Furthermore, we show that NLRP3 bound Rab11-FIP2 via its KMKK motif and that Rab11-FIP2 interacts with NLRP3 via its N-terminal C2-domain. The formation of PI4P positive endosomes and ASC-specks were also controlled by Rab11-FIP2. Collectively our results demonstrate that Rab11-FIP2 and Rab11b control NLRP3 inflammasome activation on early endosomes in human macrophages.

## Introduction

The NOD-, LRR-, and pyrin domain-containing protein 3 (NLRP3) inflammasome is a multimeric protein complex of the innate immune system that activates and controls inflammatory responses in nucleated cells (Broz & Dixit, 2016; Kelley *et al*, 2019; Lamkanfi & Dixit, 2012; Rathinam *et al*, 2012). This complex induces pyroptotic cell death and is crucial to the microbial host defense. Canonical activation of the NLRP3 inflammasome is a two-step process which initiates the processing and release of IL-1β and IL-18 that do not carry a signal peptide. First, pattern associated molecular patterns (PAMPs) unique to groups of microbes are recognized by pattern recognition receptors (PRR) to induce inflammatory signaling stimulating production of NLRP3 and pro-Interleukin-1β (pro-IL-1β) in a process referred to as priming. Lipopolysaccharide (LPS) derived from Gram-negative bacteria is frequently used to prime the NLRP3 inflammasome. The second signal is provided by a wide array of exogenous and endogenous stimuli, provided by microbial toxins or infections causing cellular damage. This leads to mitochondrial dysfunction and production of reactive oxygen species, which may cause lysosomal membrane permeability, lysosomal dysfunction and autophagic failure (Zhang *et al*, 2016).

The resulting membrane damage causes ion fluxes that trigger the NLRP3 inflammasome. Nigericin is a potent NLRP3 inflammasome activator that trigger K^+^ efflux (Munoz-Planillo *et al*, 2013). NLRP3 oligomerization drives inflammasome activation by binding the apoptosis-associated speck-like protein containing a C-terminal caspase recruitment domain (ASC), to allow ASC-speck formation (Feske *et al*, 2015; Lamkanfi & Dixit, 2014; Martin *et al*, 2014; Stehlik *et al*, 2003). The ASC-speck serves as a platform for caspase-1 mediated cleavage of pro-IL-1β to mature IL-1β, but is dispensable for caspase-1 mediated cleavage of gasdermin-D (GSDMD) and pyroptosis induction (Dick *et al*, 2016; He *et al*, 2015).

An important step in NLRP3 activation is translocation to the correct subcellular compartment for the assembly of a functional inflammasome complex. In macrophages, NLRP3 has been reported to localize to mitochondria-associated endoplasmic reticulum membranes (MAMs) (Subramanian *et al*, 2013; Zhou *et al*, 2011), the microtubule organizing center (Li *et al*, 2017) and endosomes (Zhang *et al*, 2023). Others have reported cytosolic activation complexes (Wang *et al*, 2013). Recently, NLRP3 was shown to translocate to the *trans*-Golgi network (TGN) and onto dispersed TGN (dTGN) structures containing phosphatidylinositol 4-phosphate (PI4P) following treatment with NLRP3 inflammasome inducers such as nigericin (Chen & Chen, 2018; Zhang *et al*., 2023). It was proposed that the dTGN could serve as a scaffold for NLRP3 aggregation and ASC-speck formation through ionic bonding between a conserved polybasic region in NLRP3 and the negatively charged PI4P. Now, the NLRP3 positive dTGN have been shown to be Rab5 and EEA1 positive and thus early endosomes (Chen & Chen, 2018; Zhang *et al*., 2023). It was further shown that the accumulation of 4P and the TGN resident proteins was a result of a block in retrograde trafficking caused by compounds such as nigericin (Zhang *et al*., 2023).

Rab11a and Rab11b are small GTPases which are master regulators of membrane trafficking and have redundant and overlapping functions (Joseph *et al*, 2023; Lindsay & McCaffrey, 2004). They are also important for compartmentalization of early endosomes for efficient transport to the TGN (Wilcke *et al*, 2000). Active GTP-bound Rab11 recruits effector proteins such as the Rab11-family interacting proteins (FIPs) to mediate function (Schafer *et al*, 2014). We have shown that Rab11 and Rab11-FIP2, from now on FIP2, control endosomal LPS signaling (Husebye *et al*, 2010; Klein *et al*, 2015; Skjesol *et al*, 2019; Yurchenko *et al*, 2018). We and others have shown that Rab11 and Rab11-FIP2 locate to early endosomes (Klein et al, 2015; Lindsay & McCaffrey, 2004; Skjesol et al, 2019; Zerial & McBride, 2001). Like NLRP3, FIP2 binds PI species including PI4P and locates to the TLR4 and Rab11 positive peri-nuclear endocytic recycling compartment (ERC) (Husebye *et al*, 2006; Lindsay & McCaffrey, 2004; Skjesol *et al*., 2019). In this study, we identified FIP2 as an NLRP3 binding partner that controls intracellular trafficking of NLRP3 promoting inflammasome assembly and activation. FIP2 and its binding partner Rab11b were found to coordinate caspase-1-mediated cleavage of pro-IL-1β and GSDMD, and pyroptotic cell death in human macrophages. Together, our data demonstrate that Rab11b and FIP2 control intracellular trafficking and membrane docking of NLRP3 at the PI4P positive early endosome, key events in inflammasome assembly and activation.

## Results

### FIP2 depletion strongly reduces canonical NLRP3 activation

To investigate a role of FIP2 in NLRP3 activation and pyroptosis, THP-1 cells and human macrophages were depleted for FIP2 by siRNA treatment before LPS priming and nigericin treatment. Cell death was measured by lactate dehydrogenase (LDH) release and IL-1β release by ELISA. In addition, caspase-1 mediated cleavage of pro-IL-1β into IL-1β and GSDMD into the p31 pore forming fragment were measured in cell lysates by immunoblotting. THP-1 cells silenced for FIP2 showed a 50% reduction in cell death (Fig. 1A). We next investigated if FIP2 silencing could reduce IL-1β secretion. Indeed, IL-1β release was reduced by more than 70 % as quantified by ELISA (Fig. 1B). Immunoblotting demonstrated that FIP2 silencing markedly reduced caspase-1 autocleavage giving caspase-1 p20, caspase-1 mediated cleavage of pro-IL-1β to IL-1β p17 and GSDMD to the p31 pore forming form (Fig.1C). Next, we investigated if FIP2 silencing could have a similar effect in primary human macrophages. The effect of FIP2 deletion on nigericin-induced cell death was less pronounced in primary cells than THP-1 cells but remained significant. (Fig. 1D). All donors showed a more than 70 % reduction in IL-1 β secretion as measured by ELISA (Fig. 1E). Again, immunoblotting of IL-1β confirmed the results obtained by ELISA. As in the FIP2 silenced THP-1 cells there was a marked reduction in GSDMD p31 (Fig. 1F). To further verify FIP2 as a regulator of NLRP3 inflammasome activation, THP-1 cells stably co-expressing lentiviral Flag-FIP2 (pLVX-FIP2) or Flag-Empty (pLVX-Empty) vectors were made. Following LPS priming and nigericin treatment, the cells expressing Flag-FIP2 showed a more than 100% increase in cell death and IL-1β secretion, when compared with the cells expressing Flag-Empty (Figs. 1G and 1 H).

**Figure 1.**
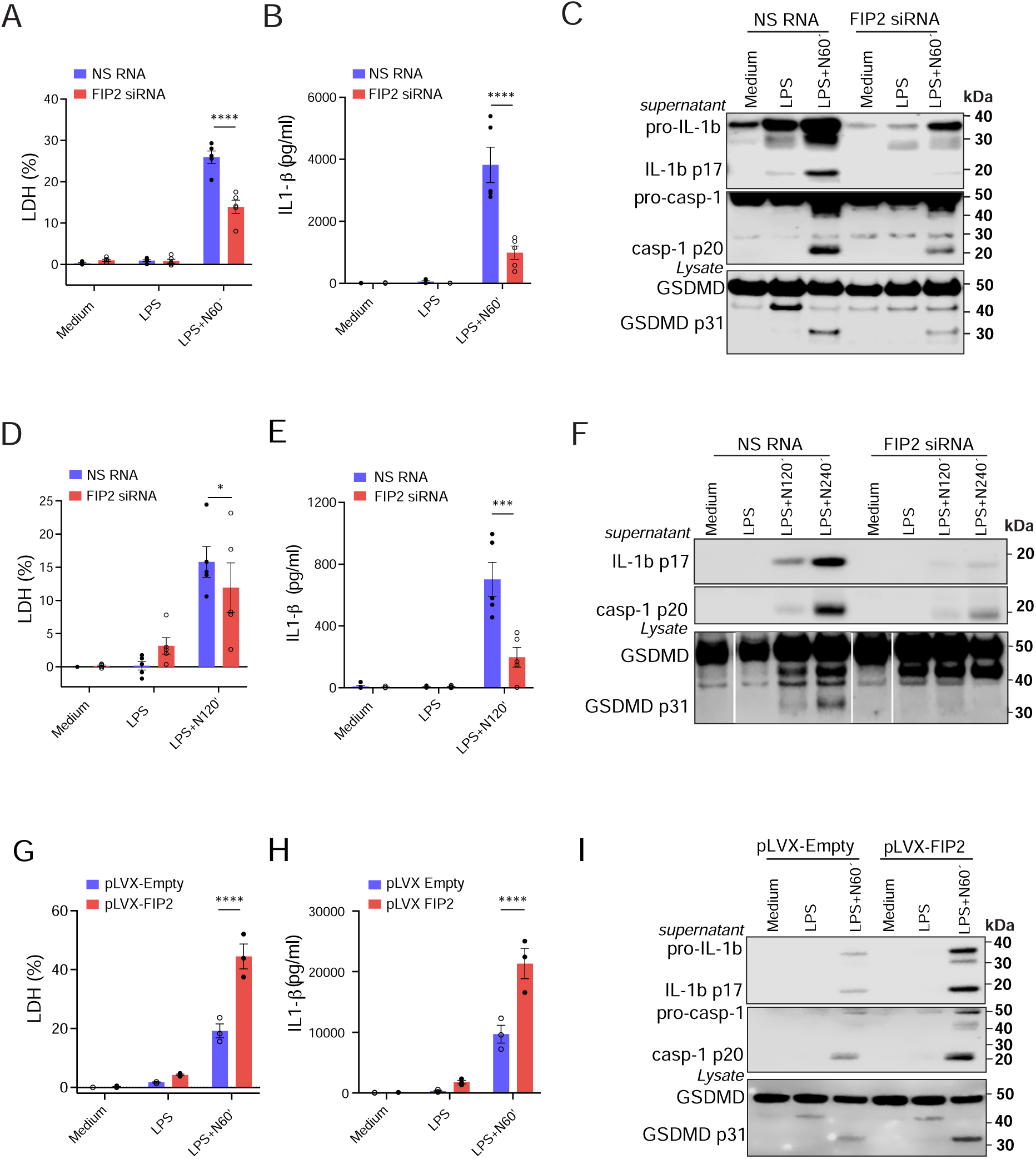
FIP2 silencing blocks pyroptosis. (**A**) Quantification of LDH-release in FIP2 silenced THP-1 cells, n = 5 independent experiments. (**B**) Quantification of IL-1β release by ELISA in FIP2 silenced THP-1 cells, n = 5 independent experiments. (**C**) Immunoblot of pro-IL-1β, IL-1β p17, pro-caspase-1, and caspase-1 p20 in FIP2 silenced THP-1 cells. (**D**) Quantification of LDH-release in FIP2 silenced human primary macrophages, n = 5 human donors. (**E**) Quantification of IL-1β release by ELISA in FIP2 silenced human primary macrophages, n = 5 human donors. (**F**) Immunoblot of IL-1β p17, caspase-1 p20, GSDMD and GSDMD p31 in FIP2 silenced human primary human macrophages. (**G**) Quantification of LDH-release in FIP2-Flag expressing THP-1 cells, n = 4 independent experiments. (**H**) Quantification of IL-1β release by ELISA in FIP2-Flag expressing THP-1 cells, n = 4 independent experiments. (**I**) Immunoblot of pro-IL-1β, β, pro-caspase-1, GSDMD, and GSDMD p31 in FIP2-Flag expressing THP-1 cells. The cells were treated with NS RNA or FIP2 siRNA before primed with 100 ng/mL LPS for 2 h and treated with 5 μM nigericin as indicated. Data information: In (A-B, D-E and G-H), data are presented as mean +/-SEM. * p = 0.029, ** p = 0,0034 and **** p < 0.0001 (Two-way ANOVA Tukey’s multiple comparisons test with adjusted p values to calculate statistical significance). N=Nigericin.

In line with these results, a marked increase in pro-IL-1β, IL-1β p17 and caspase-1 p20 was detected by immunoblotting (Fig. 1I). Cells expressing pLVX-FIP2 also showed elevated GSDMD p31 (Fig. 1I). When comparing cell death in THP-1 cells treated with the NLRP3 inhibitor MCC950 and cells with FIP2 depletion it was NLRP3 inhibition that had the larger effect, reducing cell death by 85% and 50 %, respectively (Fig EV1A and EV1B). Moreover, the NLRP3 inhibitor gave a reduction of 90% in IL-1β release, while FIP2 depletion a reduction of 75% (Figs. 1 and EV1B). Having established a role of FIP2 in NLRP3 inflammasome activation we investigated the involvement of the other type I FIPs, Rab11FIP1 (FIP1) and Rab11FIP5 (FIP5). Primary macrophages were silenced for FIP1, FIP2 or FIP5 before measuring cell death. Only the FIP2 depleted cells showed reduced cell death (Fig. EV1C). In contrast, the FIP1 depleted cells showed a modest increase, while the FIP5 depleted cells showed no change. We next monitored the presence of IL-1β p17 and caspase-1 p20 in the supernatants from the cells above by immunoblotting. The FIP5 depleted cells showed a reduction in IL-1β p17 but no change in caspase1 p20, while in the FIP1 depleted cells showed no change in either (Figs. EV1D and EV1E). The FIP2 depleted cell were the only ones that showed both reduced IL-1β p17 and caspase-1 p20. These data demonstrate that FIP2 is the only of the type I FIPs that controls both cell death and IL-1β release.

### Rab11b controls NLRP3 stimulated cell death and IL-1β secretion

Having established the involvement of FIP2 in NLRP3 inflammasome activation we next investigated the role of FIP2 binding partners, Rab11a and Rab11b. Like the FIP2 silenced cells, the Rab11b silenced THP-1 cells showed a more than 50% reduction in cell death (Fig. 2A). Quantification of IL-1β release by ELISA showed a reduction close to 50 % in the Rab11b silenced cells while the Rab11a silenced cells showed no significant change (Fig. 2B). When investigating the levels of IL-1β p17 by Western blotting, the Rab11a silenced cells showed markedly increased IL-1β p17 while in the Rab11b silenced cells it was hardly detectable (Fig. 2C). Comparing the results from the ELISA and immunoblotting, IL-1β seemed to be underscored by the ELISA, as it possibly also detects pro-IL-1β. As seen with IL-1β p17, caspase-1 p20 was also increased in Rab11a silenced cells and hardly detectable in Rab11b depleted cells (Fig. 2C). In line with these results Rab11a depletion also gave elevated GSDMD p31 while Rab11b depletion gave a marked reduction. Similar results were observed in primary human macrophages where Rab11a depletion resulted in higher IL-1β p17 and caspase-1 p20.

**Figure 2.**
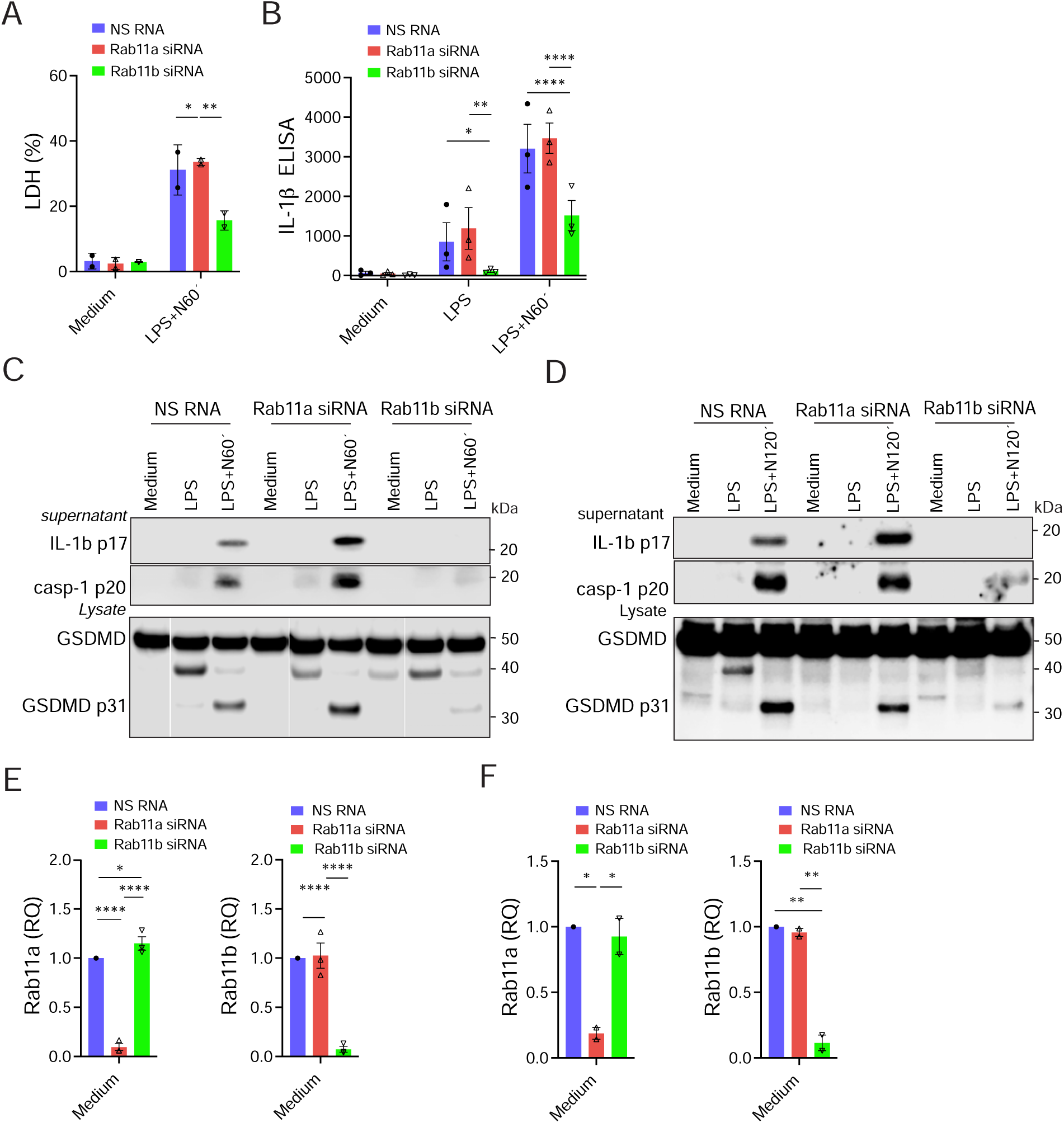
Rab11b controls both cell death and IL-1β secretion (**A**) Quantification of LDH-release in Rab11a or Rab11b silenced THP-1 cells stimulated as indicated, n = 2 experiment. (**B**) Quantification of IL-1β release by ELISA in Rab11a or Rab11b silenced THP-1 cells in supernatants, in one representative experiment of n = 3 independent experiments. (**C**) Immunoblot of IL-1β p17and caspase-1 p20 in supernatants from Rab11a and Rab11b silenced THP-1 cells; and GSDMD and β-tubulin in lysates from the same experiment, one representative experiment of n = 3. (**D**) Immunoblot of IL-1β p17 and caspase-1 p20 in supernatants from Rab11a and Rab11b silenced primary human macrophages; and GSDMD in lysates from the same donor, one representative experiment of n = 2. (**E**) Quantification of Rab11a (left panel) and Rab11b (right panel) mRNA levels by Q-PCR in Rab11a or Rab11b silenced THP-1 cells, n = 3 experiments. (**F**) Quantification of Rab11a (left panel) and Rab11b (right panel) mRNA levels by Q-PCR in Rab11a or Rab11b silenced primary human macrophages, n = 2 experiments. The cells were treated with NS RNA, Rab11a siRNA or Rab11b siRNA before primed with 100 ng/mL LPS and 5 μM nigericin as indicated. Data information: In (A and E) data are presented as mean +/-SEM and (B and F) as mean +/-SD * p = 0.012-0.0326, ** p = 0.0030-0,0061, **** p < 0.0001 (Two-way ANOVA Tukey’s multiple comparisons test with adjusted p values). N=Nigericin.

However, GSDMD p31 was slightly reduced (Fig. 2D). Rab11b depleted macrophages on the other hand showed a dramatic reduction in GSDMD p31. The efficiency of Rab11a and Rab11b silencing were verified by q-PCR and showed an equal 95% silencing efficiency (Fig. 2E). The primary human macrophages displayed a mRNA knock down efficiency of 80 % for Rab11a and 89% for Rab11b, respectively (Fig. 2F). Together, these results show that depletion of Rab11b mimics the effect of FIP2 deletion on nigericin-stimulated NLRP3 inflammasome activation in LPS primed cells an effect partly opposite to Rab11a depletion.

### FIP2 is located to the core of the ASC-specks and is needed for formation

As FIP2 controlled NLRP3 inflammasome activation, we next investigated if FIP2 could regulate ASC-speck formation. LPS primed THP-1 cells were treated with nigericin as indicated and co-stained with ASC and FIP2 or ASC and NLRP3. The spot function of IMARIS 8.2 was used to define the ASC-speck in Z-stacks from fixed cells immunostained for ASC and FIP2 or NLRP3. ASC-deficient THP-1 cells were used for correct thresholding and were verified to not secrete IL-1β (Fig. EV3B). Both FIP2 and NLRP3 were found to localize to the core of the ASC-speck in THP-1 cells (Figs. 3A and 3B). This was also seen in primary human macrophages (Figs. 3C and 3D). Quantification of ASC-specks showed that there were significantly fewer ASC-specks in FIP2 silenced THP-1 cells compared to the NS RNA treated cells (Fig. 3E). In human macrophages, ASC-speck formation needed a longer time of nigericin treatment, but also here the FIP2 silenced cells showed significantly fewer ASC-specks (Fig. 3F). The amount of NLRP3 on the ASC-specks was also quantified (Fig. EV3C). In contrast to the number of ASC-specks formed, the amount of NLRP3 on the ASC-specks did not differ between FIP2 silenced and NS RNA treated cells. Together these results demonstrate that FIP2 is located to the ASC-speck and is required for formation of the ASC-speck.

**Figure 3.**
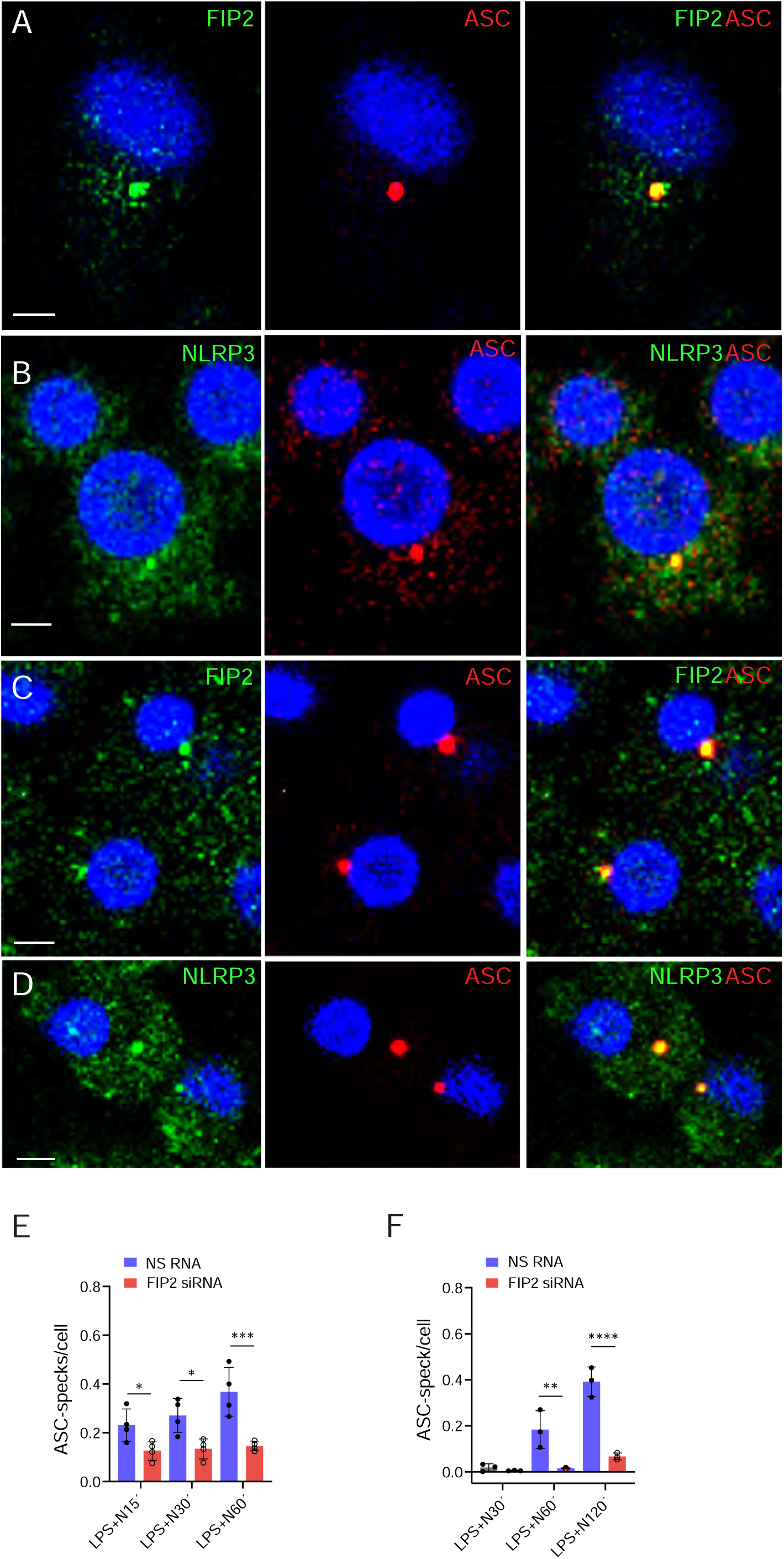
FIP2 co-localizes with ASC in the ASC speck. (**A**) Confocal image showing FIP2 (green) and ASC-speck (red). (**B**) Confocal image showing NLRP3 (green) on ASC-speck (red). The THP-1 cells were treated with NS RNA or FIP2 siRNA before LPS primed for 2 h before treatment with nigericin for 30 min (A and B). (**C**) Confocal image showing FIP2 (green) and ASC-speck (red). (**D**) Confocal image showing NLRP3 (green) and ASC-speck (red). (**E**) Quantification of ASC-speck formation in FIP2 silenced THP-1 cells, n = 4 experimental parallels, monitoring 77-230 cells per parallel. (**F**) Quantification of ASC-speck formation per cell in primary human macrophages, 242-590 cells monitored per condition. The THP-1 cells and primary macrophages were treated with NS RNA or FIP2 siRNA before primed with 100 ng/mL LPS for 2 h and treated with 5 mM nigericin as indicated (C-F). ASC-specks were identified using the IMARIS 8.2 imaging software on 3-D confocal imaging data. Data information: In (D and E) data are presented as mean +/-SD. *p=0.0112-0.0275, ** p = 0.0050, *** p = 0.0002 and **** p < 0.0001 One-way ANOVA multiple comparisons test with adj. p values. N = nigericin. Scale bar = 5 μm.

### NLRP3 binds FIP2 via its KMKK motif

Since FIP2 had a strong effect on NLRP3 inflammasome activation, we next investigated if FIP2 could bind NLRP3 and ASC. HEK293T cells were co-transfected with Flag-NLRP3 and EGFP-FIP2, with or without ECFP-Rab11. Anti-Flag-agarose-pulldowns showed that Flag-NLRP3 co-precipitated EGFP-FIP2, both in the absence and presence of Rab11 (Fig. 4A). Next, we investigated if Flag-FIP2 could co-precipitate NLRP3 and ASC. HEK293T cells were co-transfected with Flag-FIP2, ECFP-Rab11, EGFP-NLRP3 and HA-ASC.

**Figure 4.**
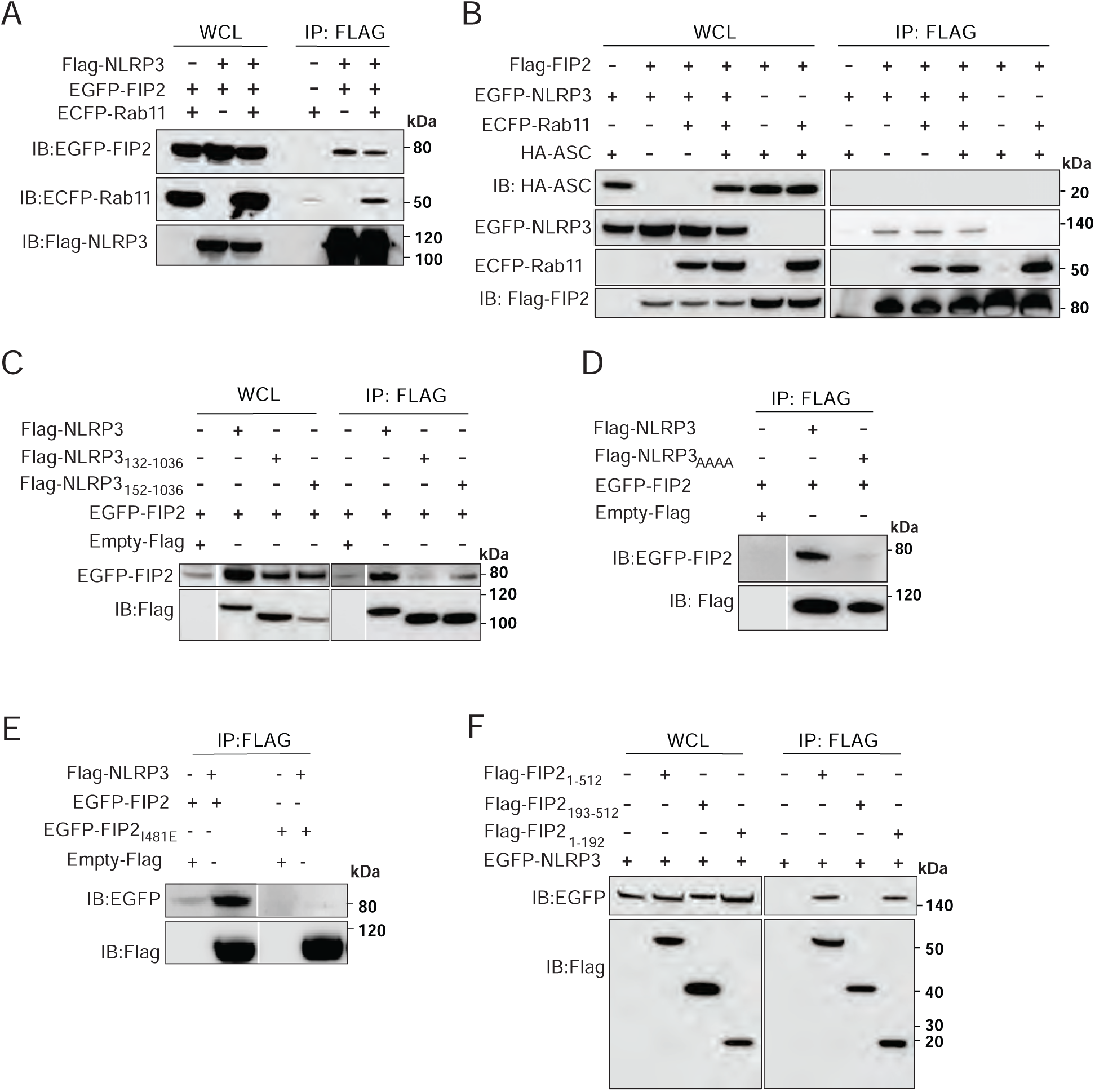
NLRP3 binds to FIP2 via its KMKK motif (**A**) Immunoblot of Flag-NLRP3 pulldown in lysates from HEK293T cells co-transfected with Flag-NLRP3, EGFP-FIP2, and ECFP-Rab11. (**B**) Immunoblot of Flag-FIP2 pulldowns in lysates from HEK293T cells co-transfected with EGFP-NLRP3, ECFP-Rab11, and HA-ASC. (**C**) Immunoblot of Flag-NLRP3, Flag-NLRP3_132-1036_ and Flag-NLRP3_152-1036_ pulldowns in lysates form HEK293T cells co-transfected with EGFP-FIP2 (**D**) Immunoblot of Flag-NLRP3 or Flag-NLRP3_AAAA_ pulldowns from HEK293T cells co-transfected with EGFP-FIP2. (**E**) Immunoblot of Flag-NLRP3 pulldowns from HEK293T cells co-transfected with EGFP-FIP2 or EGFP-FIP2_I481E_. (**F**) Immunoblot of Flag-FIP2, Flag-FIP2_1-192_ or Flag-FIP2_193-512_ pulldowns from HEK293T cells co-transfected with EGFP-NLRP3. HEK293T cells were transiently transfected with expression plasmids encoding the indicated fusion proteins for 48 h, or 24 (D – F) before immunoprecipitation by anti-Flag affinity agarose. The Empty-Flag-vector were used to ensure equal plasmid DNA amounts.

Indeed, the Flag-pulldowns showed that Flag-FIP2 co-precipitated EGFP-NLRP3 and Rab11, but not HA-ASC (Fig. 4B). It has been shown that murine NLRP3 binds PI4P through its KKKK-motif, and when mutated prevents inflammasome activation (Chen & Chen, 2018). The KKKK-motif resembles the KMKK-motif in human NLRP3 and is located at amino acids position 131 to 134. We next made a series of Flag-NLRP3 deletion mutants to locate the FIP2 binding site in the human NLRP3. The full length NLRP3 and the NLRP3 mutants, NLRP3_132-1036_ and NLRP3_152-1036_ were co-expressed with EGFP-FIP2 in HEK293T cells. The pulldown of the Flag-NLRP3 variants showed that NLRP3_132-1036_, lacking the first lysine in the KMKK motif showed no binding to EGF-FIP2. The other mutant, NLRP3_152-1036_ showed approximately 15 % of the binding observed for the wild type NLRP3 when normalized to the to the Flag intensity of the respective pulldown (Fig. 4C). To verify the involvement of NLRP3 KMKK-motif in FIP2 binding, we next made a NLRP3 mutant where the KMKK-motif was mutated to AAAA, NLRP3_AAAA_. When comparing the Flag-NLRP3 and Flag NLRP3_AAAA_ pulldowns by immunoblotting, NLRP3_AAAA_ showed almost no binding to FIP2 compared to the NLRP3 full-length (Fig. 4D). We also included FIP2I481E, a mutant defective in Rab11 binding (Jagoe *et al*, 2006). Indeed, the pulldowns of FIP2I481E, showed that Flag-NLRP3 did not bind FIP2I481E, suggesting that also Rab11 contributes to FIP2‘s binding to NLRP3 (Fig 4E). To investigate what part of FIP2 that was responsible for NLRP3 binding we next did pulldowns with full-length FIP2 and the FIP2 deletion mutants, Flag-FIP2_1-192_ and Flag-FIP2_193-512_ (Fig. 4F). Interestingly, while the N-terminal FIP2_1-192_ variant containing the C2-domain showed strong binding, the C-terminal FIP2_193-512_ deletion mutant showed no binding. Together these results show that the KMKK motif in NLRP3 is the FIP2 binding site and that the NLRP3 binding motif in FIP2 is in the N-terminal FIP2 C2-domain. Furthermore, our data suggest that NLRP3 can be found in complex with FIP2 and Rab11, and that Rab11 to FIP2 is required for interaction of FIP2 with NLRP3.

### FIP2 controls NLRP3 translocation to the TGN during LPS priming

We next investigated the cellular location of NLRP3 during LPS priming. We have previously shown that TLR4 is enriched in the peri-nuclear Rab11-positive ERC (Husebye *et al*., 2010; Klein *et al*., 2015). Recently, NLRP3 was shown to accumulate in the TGN of murine and human macrophages, and on dTGN structures following nigericin treatment (Andreeva *et al*, 2021; Chen & Chen, 2018; Nanda *et al*, 2021; Zhang *et al*., 2023). As seen in Figure 5A, there are low NLRP3 amounts in the TGN of THP-1 cells before stimulation and high amounts following LPS priming (Figs. 5A and 5B). Quantification of NLRP3 showed that the level of NLRP3 in the TGN was increased by 257% in LPS primed THP-1 cells (Fig. 5C). Then, we investigated if FIP2 could regulate NLRP3 translocation to the TGN. In the TGN of unstimulated cells the levels of NLRP3 were the same, while in FIP2 silenced cells the NLRP3 levels showed a significant a reduction of 66%. We also investigated if treatment of THP-1 cells with the NLRP3 inflammasome inhibitor, MCC950, could affect NLRP3 in the peri-nuclear TGN following LPS priming. If so, this could indicate that TGN-localized NLRP3 was in an active form or not. As seen in Figure 5D, MCC950 did not affect NLRP3 in the TGN, neither in resting nor in LPS primed cells. Also, in primary human macrophages FIP2 and NLRP3 showed higher intensities in the TGN following LPS priming. In contrast to the THP-1 cells the TGN consisted of more tubular structures occupying most of the cell (Fig. EV5). In THP-1 cells FIP2 showed a clear location to the TGN that was significantly enhanced following LPS priming (Figs. EV5E-EV5G). These results show that FIP2 controls NLRP3 translocation to the TGN-ring during LPS priming.

**Figure 5.**
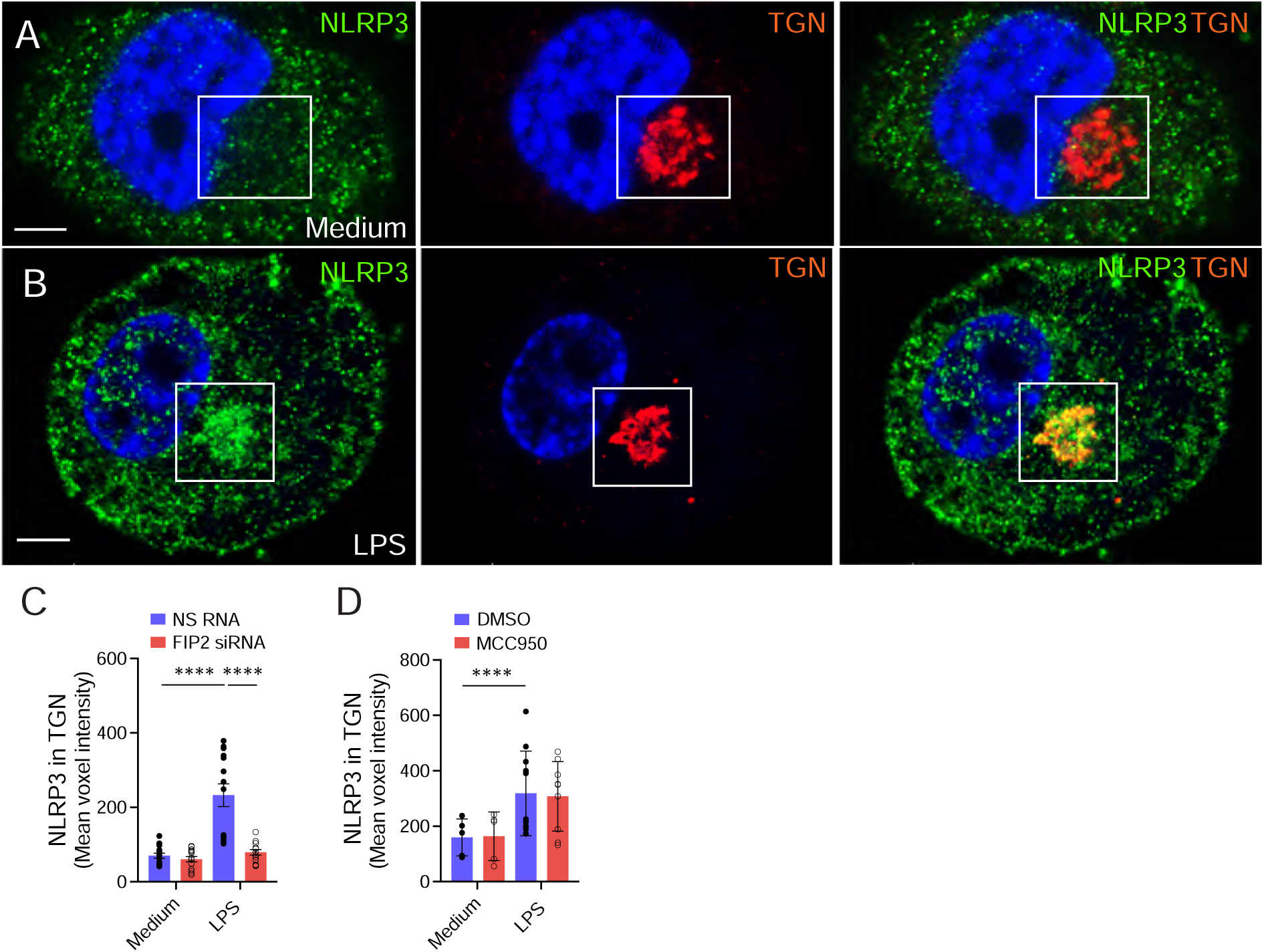
FIP2 recruits NLRP3 to the TGN during LPS priming **(A)** Confocal image showing NLRP3 (green) and TGN46 (red) in resting THP-1 cells. (**B**) Confocal image showing NLRP3 (green) and TGN46 (red) in LPS primed THP-1 cells. (**C**) Quantification of NLRP3 in the TGN-ring structure in unstimulated and LPS primed THP-1 cells, 50-226 cells were monitored in n = 4 independent experiments. (**D**) Quantification of NLRP3 in the TGN-ring structure in unstimulated and LPS stimulated THP-1 cells pretreated with DMSO or MCC950 5 min, 55-155 cells monitored in n = 2 independent experiments. The cells were left unstimulated, primed with LPS for 2 h. MCC950 were added 30 min prior LPS priming. The TGN46 positive peri-nuclear ring-structures were identified using the IMARIS 8.2 imaging software on 3-D confocal imaging raw data. Data information: In (C) data are presented as mean +/-SEM, and (D) as mean +/-SD. **** p < 0.0001 (Two-way ANOVA, Tukey’s multiple comparisons test with adj. p values). N = nigericin. Scale bar = 5 μm.

### FIP2 regulates NLRP3 recruitment to the dTGN during nigericin treatment

NLRP3 positive dTGN has been shown to serve as a scaffold for NLRP3 aggregation driving ASC-speck formation (Andreeva *et al*., 2021; Chen & Chen, 2018). Thus, we examined if FIP2 could locate to and affect the number of nigericin stimulated dTGN primary macrophages. The dTGN was identified using the spot function of IMARIS 8.2 on in multi-channel Z-stacks from fixed cells immunostained for ASC and NLRP3 or ASC and FIP2. As previously reported we found the dTGN to be strongly positive for NLRP3 (Chen & Chen, 2018) (Figs. 6A and 6C). Interestingly, dTGN structures were also strongly positive for FIP2 (Fig 6B). dTGN were largely small puncta but also varied in size up to enlarged structures with a defined limiting membrane (Figs. 6A and 6B). Next, we investigated dTGN formation and the NLRP3 and FIP2 levels in nigericin treated LPS primed THP-1 cells (Figs. 6C and 6D). Also, here the dTGN appeared largely as small puncta positive for NLRP3 or FIP2 (Figs. 6C and 6D). Interestingly, the FIP2 silenced cells showed a significant 60 %reduction of dTGN numbers and FIP2 suppression reduced NLRP3 on dTGN by 65% (Fig. 6E). The FIP2 levels on the remaining dTGN of the FIP2 silenced cells were reduced to similar levels as the negative control cells, confirming specificity of the FIP2 antibody used (Fig. 6F, right panel). The NLRP3 positive dTGN has previously been shown to be Rab5 and EEA1 positive early endosomes (Zhang *et al*., 2023).

**Figure 6.**
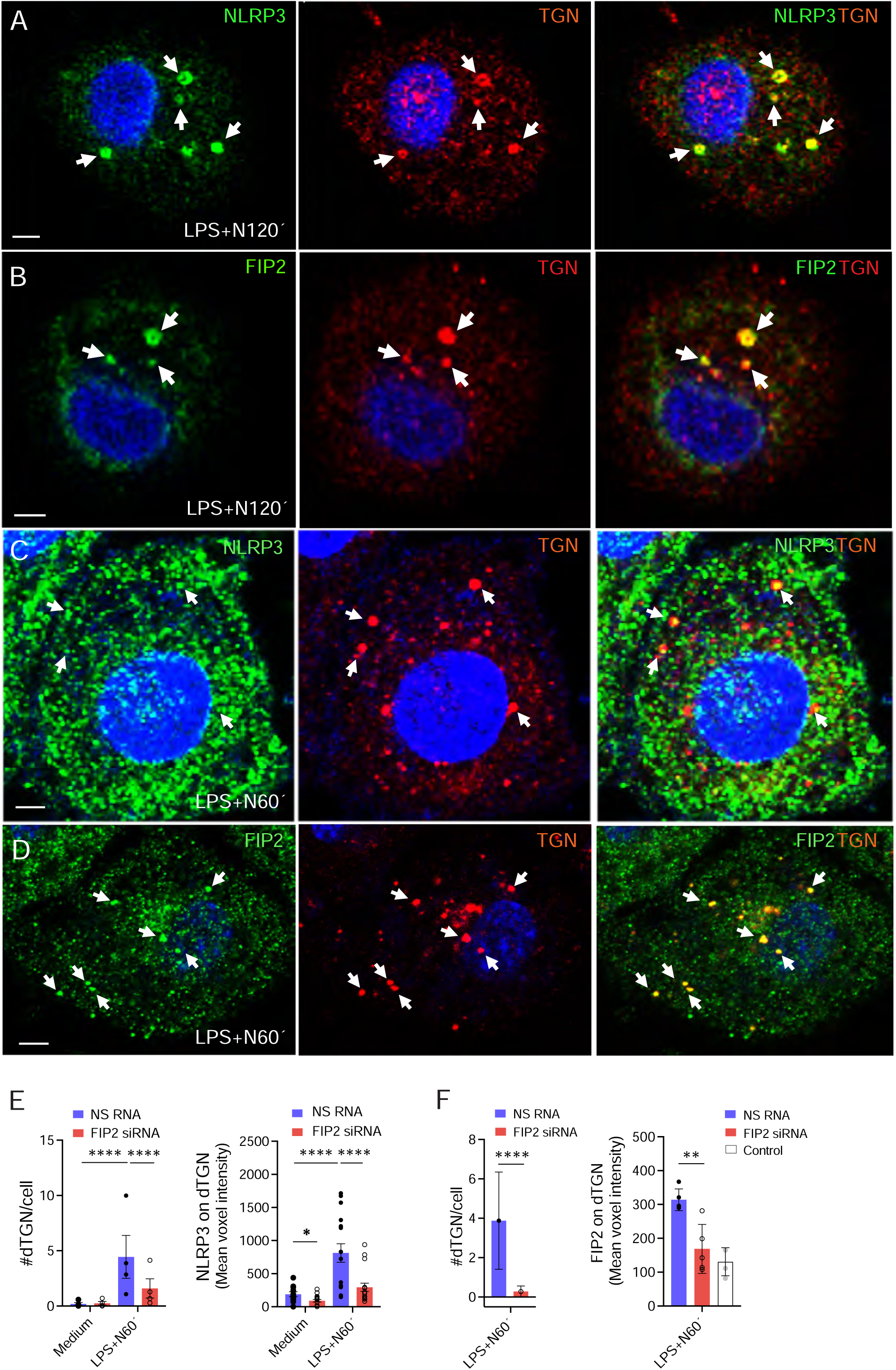
FIP2 locates to the dTGN structures and controls their number (**A**) Confocal images showing NLRP3 in TGN46 positive dTGN structures in primary human macrophages. (**B**) Confocal images showing NLRP3 in TGN46 positive dTGN structures human primary macrophages. The macrophages were LPS primed and treated with nigericin for 2 h (A and B). (**C**) Confocal images showing Flag-FIP2 in microdomains on Rab5 coated endosomes. (**D**) Confocal images showing NLRP3 microdomains co-localizing with FIP2 on a FIP2 coated enlarged endosome. (**E**) Quantification of dTGN numbers and corresponding NLRP3 levels in THP-1 cells stimulated as indicated. 96-126 cells were monitored per condition, in n = 4 independent experiments. (**F**) Quantification of dTGN numbers and corresponding NLRP3 levels in THP-1 cells stimulated as indicated. 62-132 cells were monitored per condition, in n = 1 independent experiment. THP-1 cells were unstimulated or LPS primed for 2 h before 1 h of nigericin treatment. TGN46 positive dTGN structures were identified by the IMARIS 8.2 imaging software and the NLRP3 intensities given as mean voxel intensity. Data information: In (E) data are presented as mean +/-SEM, and (F) as mean +/-SD. **** p < 0.0001 (Two-way ANOVA, Tukey’s multiple comparisons test with adj. p values). White arrows point at dTGN puncta positive for NLRP3 or FIP2; or Rab5 or FIP2 coated endosomes positive for FIP2 or NLRP3, respectively. N = nigericin. Scale bar = 5 μm.

Indeed, LPS primed THP-1 cells treated with nigericin showed EEA1 positive dTGN (Fig. EV6A). Some of these clearly displayed an enlarged phenotype with a defined limiting membrane. We also found enlarged EEA1 endosomes showing NLRP3 positive microdomains in the limiting membrane (Fig. EV6B). Together, these results support that the dTGN structures are early endosomes and that FIP2 is instrumental in their formation.

*NLRP3 and FIP2 are located on Rab5 positive endosomes during inflammasome activation* Given the role of FIP2 in controlling the number of dTGN, we investigated if FIP2 could control the PI4P positive puncta formed during NLRP3 inflammasome activation. After nigericin treatment of primed cells, PI4P was found on NLRP3 positive structures, varying in size from small puncta to endosomes with visual limiting membrane (Fig. 7A). The small NLRP3 and PI4P positive puncta were by far the most abundant. Moreover, we also observed the formation of endosomes positive for NLRP3 and Rab5, with a similar phenotype as the PI4P-positive endosomes (Fig 7B). We could also identify Rab5 endosomes with PI4P microdomains (Fig. 7C). Interestingly, we observed FIP2 positive endosomes with NLRP3 and FIP2 co-localization in microdomains (Fig. 7D). The PI4P-positive puncta, most likely small early endosomes or punctuate microdomains in the limiting membrane of enlarged endosomes. As observed for TGN46 in the limiting membrane of enlarged EEA1 endosomes (Fig. EV6A), for PI4P on Rab5 endosomes (Fig. 7C) and for NLRP3 on FIP2 or Rab11b endosomes (Figs. 7D and EV7C). In some cases, PI4P also covered larger regions of the endosome limiting membrane (Fig. 7A), as was the case for NLRP3 (Figs. 7B and EV6B). Interestingly, NLRP3 inflammasome activation increased the number of PI4P positive endosomes by 400% (Fig. 7E). FIP2 silencing reduced the number of PI4P positive endosomes by more than 70 %, a level close to the unstimulated cells (Fig. 7E, left panel). Also, we observed a significant reduction in the amount of NLRP3 on PI4P positive endosomes (Fig. 7E, right panel). We next investigated if the number of PI4P positive endosomes was affected by the NLRP3 inhibitor MCC950. If so, it would be indicative of NLRP3 being in an oligomerized active state on the PI4P endosomes. Indeed, the MCC950 treatment reduced the number of PI4P positive endosomes, and the reduction was statistically significant (Fig. 7F). Because NLRP3 activation also was dependent on Rab11b we used THP-1 cells co-expressing Flag-Rab11b to investigate if Rab11b was located on Rab5 endosomes. Indeed, Rab11b was frequently found on Rab5 enlarged endosomes following nigericin treatment of LPS primed cells (Fig. EV7). We also detected microdomains that contained Rab11b.

**Figure 7.**
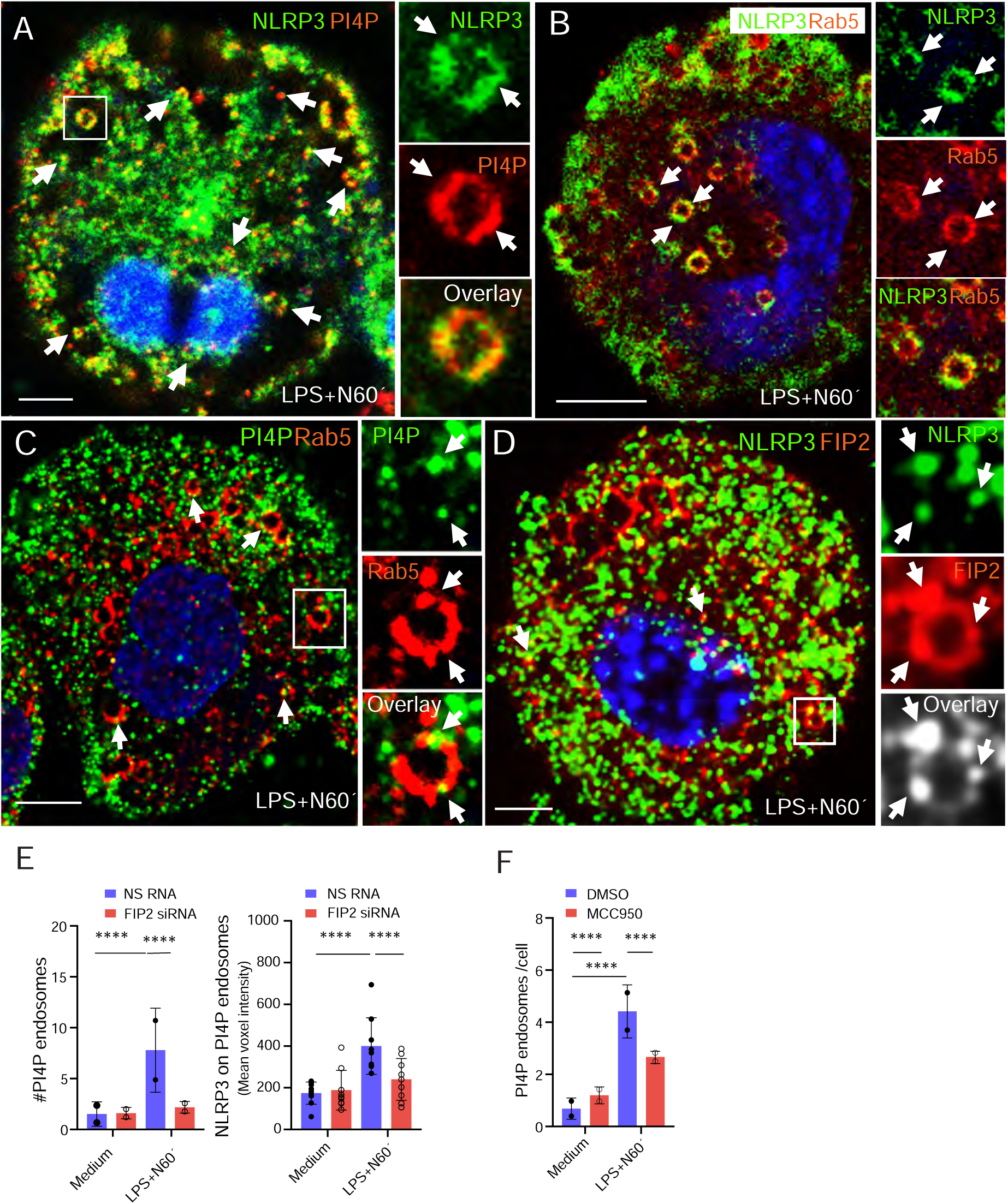
PI4P shifts from the TGN and onto endosomal like structures that enlarges following nigericin treatment. (**A**) Confocal image showing PI4P in the TGN-ring of resting cells. (**B**) Confocal image showing that PI4P is depleted form the TGN46 positive ring and appear on puncta endosomal structures following LPS priming. (**C**) Confocal image showing NLRP3 (green) on an enlarged PI4P positive endosome (red). (**D**) Confocal image showing PI4P (green) on an enlarged Rab5 coated endosome (red). (**E**) Quantification of PI4P levels in the TGN before and after LPS priming for 2 h. (**F**) Quantification of PI4P positive endosomes before and after nigericin treatment of LPS primed cells. (**G**) Quantification of PI4P positive endosomes before and after nigericin treatment of LPS primed cells. The cells were primed with 100 ng/mL LPS for 2 h and treated with 5 μM Nigericin. MCC950 were added 30 min prior LPS priming. In (E) data are presented a as mean +/-SEM. **** p<0.0001 (Two-way ANOVA Tukeýs multiple comparisons test with adj. p values) and (F) as mean +/-SD. * p=0.029 (Multiple unpaired t test). Scale bar = 5 μm.

**Figure 8.**
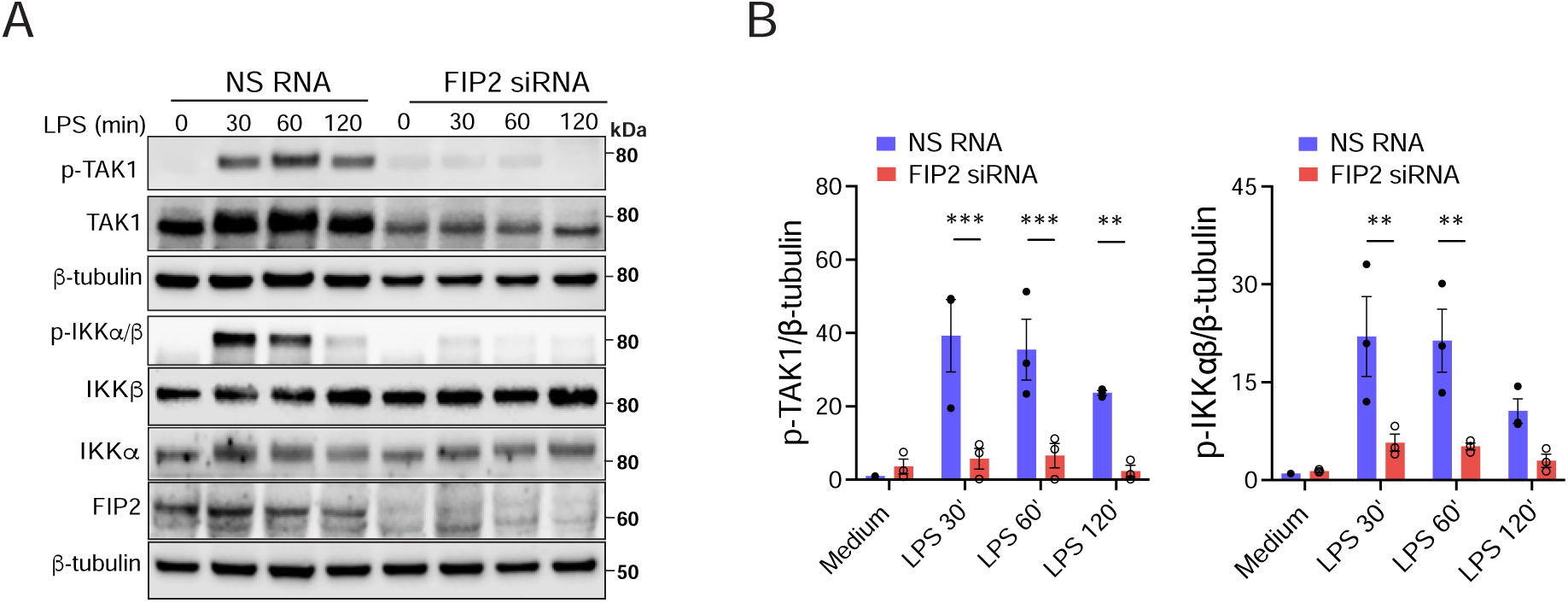
FIP2 control LPS stimulated IKKβ activation through TAK1 (**A**) Immunoblots showing LPS stimulated TAK1 phosphorylation, and IKKβ/IKKβ phosphorylation together with total levels of TAK1, IKKβ, IKKα, FIP2 and β-tubulin. NS RNA or FIP2 siRNAs treated THP-1 cells were primed with 100 ng/mL LPS as indicated. (**B**) Quantification of TAK1 phosphorylation (left panel) and IKKβ phosphorylation (right panel) normalized versus β-tubulin from n = 3 independent experiments. In (B) data are presented a as mean +/-SEM. ** p=0.0019-0.0071, *** p=0.0002-0.0007 (Two-way ANOVA Tukey’s multiple comparisons test with adj. p values).

The enlarged Rab11b endosomes also showed NLRP3 and PI4P positive microdomains. Together these results show that both FIP2 and Rab11 regulate the dynamics of NLRP3 and PI4P on early endosomes.

### FIP2 controls LPS stimulated IKKβ activation

IKKβ activation has been reported to be the cue for NLRP3 translocation to the TGN (Nanda *et al*., 2021; Schmacke *et al*, 2022). We next investigated if FIP2 could affect LPS stimulated IKKβ activation, suggesting a mechanism whereby FIP2 could drive NLRP3 translocation to the TGN. The TAK1 kinase complex phosphorylates and activates IKKβ (Wang *et al*, 2001; Zhang *et al*, 2014). Interestingly, in FIP2 silenced THP-1 cells LPS stimulated phosphorylation of TAK-1 was reduced by more than 80 % at all timepoints of LPS stimulation, while IKKβ activation measured by phosphorylation were decreased by more than 75 % at all timepoints of LPS stimulation (Figs. 6A and 6B). FIP2 depletion was verified by immunoblotting. Together, these results show that FIP2 control TAK-1 mediated LPS IKKβ activation and is also required for TAK1 stability.

## Discussion

Recruitment of NLRP3 to endosomal membranes is required for proper inflammasome activation. In this work we demonstrate that FIP2 and its binding partner, Rab11b, are critical regulators of NLRP3 location and inflammasome activation. This conclusion is based on our observations that suppression of FIP2-or Rab11b strongly inhibited caspase-1 mediated cleavage of pro-IL-1β and GSDMD. FIP2 bound to the polybasic region of NLRP3 and controlled its localization to endosomal membranes and ASC-specks. IKKβ phosphorylation, a priming step that recruits NLRP3 to TGN structures (Schmacke *et al*., 2022), was strongly reduced by FIP2 depletion which may explain the potent regulatory effects of FIP2 on NLRP3 recruitment to endosomal membranes and subsequent inflammasome activation. NLRP3 is by far the most studied inflammasome and is regulated by a diversity of factors (Swanson *et al*, 2019). Here we found that Rab11b and FIP2 were essential regulators of the canonical NLRP3 inflammasome activation in human macrophages. Interestingly, Rab11b but not Rab11a, controlled NLRP3 inflammasome activation describing a new role of Rab11b and FIP2. We have previously shown that Rab11a controls phagocytosis of *E. coli* and subsequent production of the type I interferon IFN-β (Husebye *et al*., 2010). Thus, Rab11a and Rab11b may have differential roles in regulating innate immune responses in human macrophages.

The conserved KKKK motif in the polybasic region in NLRP3 in mice was recently shown to bind the negatively charged PI4P enriched on early endosomes (Chen & Chen, 2018).

The motif serves as a PI4P-binding domain for NLRP3 recruitment and activation in mouse macrophages (Chen & Chen, 2018). In human NLRP3, the second lysine in the KKKK motif is replaced with methionine, giving KMKK. Surprisingly, we found that the deletion mutant of human NLRP3 missing the first lysine at amino acid position 132 was deficient in FIP2 binding. Furthermore, when the KMKK motif was substituted with AAAA, FIP2 binding was lost. Thus, NLRP3 binding to FIP2 clearly depends on the KMKK motif. Also, Rab11 binding to FIP2 seemed to be involved since a substitution in the Rab11 binding motif resulted in loss of interaction between FIP2 and NLRP3.

The translocation of NLRP3 to the TGN has been reported to be dependent on IKKβ activation, and IKKβ is required for the assembly and activation of the NLRP3 inflammasome (Nanda *et al*., 2021; Schmacke *et al*., 2022). We here found that the levels of NLRP3 and FIP2 in the perinuclear TGN-were markedly increased by LPS priming (Figs. 5 and EV5). IKKβ binds NLRP3 and drives NEK7 independent priming in human macrophages by recruiting NLRP3 to PI4P (Asare *et al*, 2022; Schmacke *et al*., 2022). TAK1 is an important regulator of NLRP3 inflammasome activation (Okada *et al*, 2014), and TAK1 phosphorylates IKKβ to activate it (Zhang *et al*., 2014). In this study, we showed that FIP2 depletion restricted LPS stimulated TAK1 phosphorylation, IKKβ activation and subsequent NLRP3 inflammasome activation and cell death. Thus, we have demonstrated that FIP2 translocate NLRP3 to the TGN and onto endosomal membranes by a mechanism involving TAK1 mediated phosphorylation of IKKβ. NLRP3 activators such as nigericin cause K^+^ efflux and disrupt endosome-TGN retrograde transport resulting in accumulation of *trans*-Golgi markers and PI4P on endosomes (Chen & Chen, 2018; Lee *et al*, 2023; Zhang *et al*., 2023). The dTGN structures are known to be early endosomes (Zhang *et al*., 2023), an observation that was confirmed by us in this report. Inhibition of retrograde transport causes accumulation of proteins on endosomes as well as endosomal enlargement (Huotari & Helenius, 2011).

Our results show that FIP2 depletion markedly reduced ASC-speck formation as well as NLRP3 and PI4P levels on early endosomes, pointing to an important role of FIP2. We also found that nigericin treatment of LPS primed cells resulted in considerable enlargement of NLRP3-and PI4P positive endosomes (Fig. 7, and Figs. EV7A and EV7C).

Interestingly, these enlarged endosomes showed microdomains of both PI4P and NLRP3 as well as FIP2 and NLRP3 co-localization on these endosomes (Fig. 7D). The presence of FIP2 on NLRP3 positive microdomains could point to a mechanism where FIP2 is involved at the endosomal hub for ASC-speck formation. Cells co-expressing Flag-Rab11b frequently showed enlarged early endosomes positive for NLRP3 and PI4P and increased levels of NLRP3 activation measured as caspase-1 mediated cleavage of GSDMD (Fig. EV7). Concomitantly, increased GSDMD cleavage was observed in inflammasome activated cells. This also supports that NLRP3 inflammasome activation occurs from enlarged endosomes.

It is known that PI4-Kinases like PI4KIIα generates endosomal PI4P that regulates receptor sorting at early endosomes (Henmi *et al*, 2016). Early endosome-to-TGN transport is organized by the retromer complex (Wassmer *et al*, 2009). FIP2 interacts with EHD3, a PI4P binding protein, that regulates retrograde transport from the early endosome (Naslavsky *et al*, 2006). Interestingly, also actin-polymerization is required for retromer function (Hao *et al*, 2013). Others have shown that FIP2 regulates actin-cytoskeleton dynamics (Dong *et al*, 2016), and we have shown that Rac1 and CdC42 stability depends on FIP2 in human macrophages (Skjesol *et al*., 2019). Also, FIP2‘s partner Rab11, regulates compartmentalization of early endosomes and is required for retrograde transport (Wilcke et al, 2000). So, is therefore likely that FIP2 also may control retromer function.

In summary, we have uncovered a novel role for FIP2 in controlling NLRP3 inflammasome activation in a Rab11b specific manner. We have found that FIP2 interacted with NLRP3 and performed intracellular trafficking that resulted in localization of NLRP3 to endosomal membranes with subsequent inflammasome activation. These data implicate that coordination of vesicle traffic by Rab11b and its effector molecule FIP2 play an essential role in antimicrobial defense and sterile inflammatory diseases.

## Materials and methods

### Reagents

K12 LPS (tlrl-eklps) and nigericin (tlrl-nig) were from Invivogen. The following primary antibodies were used mouse anti-Caspase-1 (p20) (Bally-1) (AG-20B-0048-C100) and rabbit-anti ASC (AL177) (AG-25B0006F-C100). Mouse anti IL-1 beta/IL-1F2 (MAB201) and goat anti-mouse IL-1β (AF-401-NA) from R&D systems. Rabbit anti-Rab11FIP2 [EPR12294-85] (ab180504), rabbit anti-Rab11FIP2 (ab174313), rabbit anti-β-tubulin (ab6046) and goat-anti-NLRP3 (ab4207), rabbit anti-TGN38 (ab50595) were from Abcam. Rabbit anti-GFP (632592) and mouse anti-GFP a (632381) were from Takara Bio. Mouse anti–FLAG M2 was from Sigma-Aldrich. Anti goat-FIP2 (S-17) (sc-163274) and mouse anti-TGN38/46 (sc-166594) were from Santa Cruz Biotech. Rabbit anti-Gasdermin D (#96458), rabbit anti-Gasdermin D (#97558), rabbit anti-phospho-TAK1 (#4508), rabbit anti-phospho-IKKα/IKKβ (16A6) (#2697), rabbit anti-IKKα (#2682) and rabbit anti-IKKβ (D30C6) (#8943), rabbit anti-Rab5 (C8B1) (#3547) were from Cell Signaling Technology. Rabbit anti-IL-1 β (Cleaved Asp116) (PA-105048), rabbit anti-NLRP3 (SC06-23), rabbit anti-NLRP3 (MA5-32255), rabbit anti-TGN46 (PA5-23068), mouse anti-Rab5 a (66339-1-IG), rabbit anti-Rab5 y (PA5-29022) and rabbit anti-EEA1 (PA5-17228) from Thermo Fisher Scientific. Purified Anti-PI4P IgM (Z-P004), was from Echelon Biosciences. Secondary antibodies used for imaging: Chicken anti-rabbit IgG Alexa Fluor 488 (A-21443), chicken anti-rabbit IgG Alexa Fluor 647 (A-21443), donkey anti-goat IgG (H+L) Alexa Fluor Plus 488 (A-32814), donkey anti-goat IgG (H+L) Alexa Fluor Plus 647 (A-32849), donkey anti-mouse IgG (H+L) Alexa Fluor Plus 488 (A32766), donkey anti-mouse IgG (H+L) Alexa Fluor Plus 647 (A32787), rabbit anti-mouse IgG Alexa Flour 488 (A-11059), rabbit anti-mouse IgG Alexa Flour 647 (A-21239), rabbit anti-mouse IgG Alexa Flour 647, goat anti-mouse IgM (A-21042) and rabbit anti-mouse IgM FITC (31557) were from Thermo Fisher Scientific.

### Cells and cell lines

THP-1 cells (monocytic cell line derived from a patient with acute monocytic leukemia ATCC®, TIB-202™) or ASC-deficient THP-1 (Invivogen, thp-dasc) were cultured in RPMI-1640 medium (Sigma, R8758) supplemented with 10 % heat-inactivated fetal bovine serum (FBS) (Gibco, 10270106), 700 μM L-glutamine (Sigma-Aldrich, Merck G7513) and 5 μM 2-mercaptoethanol (Thermo Fisher Scientific, Gibco™ 31350-010) at 37 °C and 5 % CO_2_.

Differentiation into macrophage-like THP-1 cells was induced by 60 ng/mL phorbol 12-myristate 13-acetate (PMA) (Sigma-Aldrich, P8139). Human peripheral blood mononuclear cells (PBMCs) were isolated from buffy coats from the Blood Bank at St. Olav‘s Hospital (Department of Immunology and Transfusion Medicine) by density gradient centrifugation using Lymphoprep^TM^ (Axis-Shield, AXS-1114547). Monocytes were isolated by adherence culture plates in RPMI-1640 supplemented with 5% pooled A+ serum (Department of Immunology and Transfusion Medicine, St. Olav‘s Hospital) at 37 °C and 5% CO_2_ for 45 min before washing three times in Hank‘s Balanced Salt solution (Sigma-Aldrich, H9269). Monocytes were further differentiated into macrophages for 10 days in RPMI-1640 supplemented with 10% A^+^ serum (Department of Laboratory Medicine, Unit of Immunology and Transfusion Medicine, St Olavs University Hospital), 700 μM L-glutamine, 100 nM penicillin/streptomycin and 25 ng/mL recombinant human M-CSF/CSF1 (R&D systems, 216-MC-025) with a medium change on day 3. HEK293T cells (ATTC, CRL-11268™) were cultured in DMEM (BioWhittaker®, 12-604F) supplemented with 10% FBS, 700 μM L-glutamine and 100 nM penicillin/streptomycin at 37 °C and 8% 1 μg/mL of Ciprofloxacin Hydrochloride (Corning cellgro™ 61-277-RG) and 0.5 mg/mL G418 (Geneticin™) (Thermo Fisher Scientific, Gibco™ 10131035). All cell lines were regularly checked for mycoplasma contamination.

### siRNA treatment

THP-1 cells were transfected with 16 nM siRNA in LipofectamineÔ RNAiMax (Thermo Fisher Scientific, 13778075) 24 h after seeding in media without antibiotics. Fresh media (without PMA) was added after 48 h, and cells were stimulated after 48 hours of rest. Primary macrophages were transfected with 20 nM siRNA on day 6 and 8 after seeding, using LipofectamineÔ 3000 (Thermo Fisher Scientific, L3000001). The AllStars Negative Control siRNA (QIAGEN, SI03650318) was used as a non-silencing control and termed NS RNA. Hs_RAB11FIP1_12 (SI02778972), Hs_RAB11FIP2_5 (SI04305672), Hs_RAB11FIP5_5 (SI03246782), Hs_RAB11A_5 FlexiTube siRNA (SI04305672) and Hs_RAB11B_6 FlexiTube siRNA (SI02662695) were all from Qiagen and used to target FIP1, FIP2 and FIP5 respectively.

*Generation of a stable THP-1 cell line co-expressing Flag-FIP2, Flag-Rab11a or Flag-Rab11b* THP-1 expressing lentiviral encoding FIP2 was made by cloning FIP2 into the bicistronic lentiviral expression vector pLVX-EF1α-IRES-Puro-Vector (Takara, 631988) and co-transfected with packaging plasmids psPAX2 and pMD2.G, kindly provided by the TronoLab (Addgene plasmids 12260 and 12259, to produce pseudo viral particles in HEK293T cells (ATTC CRL-3216™). The pEGFP-FIP2 (KIAA0941 sequence in pEGFP-C1) was used as a template for FIP2. Supernatants were collected at 48 h and 72 h, combined and concentrated using Lenti-X™ Concentrator (Takara, 631232). Subsequently, the viral particles were used for transduction of THP-1 wild-type cells and successful transduction of Flag-FIP2 verified by Western blotting.

### Cloning of Flag NLRP3 full length and NLRP3 deletion mutants

Phusion High-Fidelity DNA Polymerase (Thermo Fisher Scientific) was used for amplification of NLRP3 gene sequences using the pEGFP-C2-NLRP3 plasmid as template for making the Flag-NLRP3 variants. The pEGFP-C2-NLRP3 was a gift from Christian Stehlik (Addgene plasmid # 73955; http://n2t.net/addgene:73955; RRID:Addgene_73955). PCR products, or restricted vectors, were purified by QIAquick PCR purification and gel extraction kits (Qiagen, 28704). Endofree plasmid Maxi kit (Qiagen, 12362) was used for endotoxin-free plasmids preparations. The PCR products were subcloned into pLVX-EF1α-IRES-Puro vector (Takara Bio, 631988) Sequencing of plasmids was done at Eurofins Genomics.

Primers used for cloning:

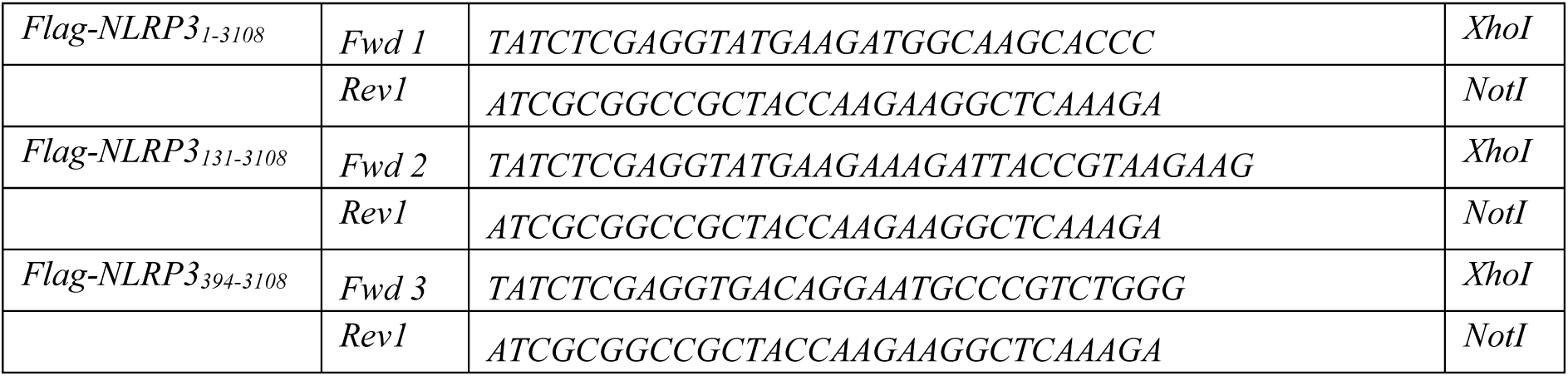

### Cloning of the NLRP3_AAAA_ mutant

Two PCR products were prepared using the full length Flag-NLRP3_1-3108_ plasmid as a template and following primers:

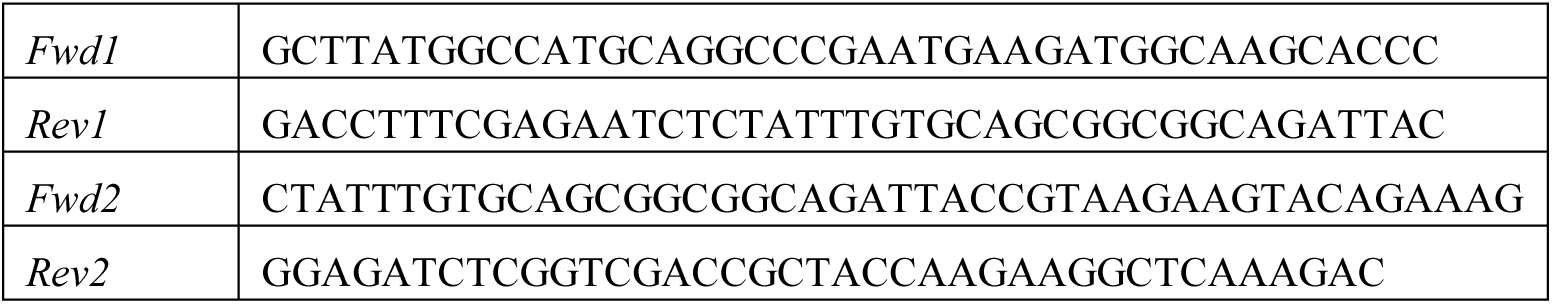

The PCR products were purified from agarose after being separated by agarose gel electrophoresis (QIAquick PCR Purification Kit, 28106) and combined with cleaved vector for combining DNA fragments using a Gibson Assembly kit (Thermo Fisher Scientific, A46624) following manufacturer recommendations for three DNA fragments. The kit was used to combine the EcoRI-HF (New England Biolabs, R3101S) cleaved empty pCMV-DYKDDDDK-N vector (Takara Bio, 635688) with 2 PCR products to change the NLRP3 KMKK motif to AAAA. The final ligation mix was transformed into competent *E. coli* DH5α. The final NLRP3_AAAA_ plasmid vector was verified by sequencing (Eurofin Genomics).

### LDH assay

Cell culture supernatants were collected and centrifuged at 10 000 x g for 1 min followed by LDH release analysis using LDH Cytotoxicity Assay Kit (Cayman Chemical, 601170) following the manufacturer’s instructions. On the day of the experiment medium was changed into medium containing 1 % FBS or 1 % human A^+^ serum. This to allow better reability of the LDH assay. Following stimulations with 100 ng/mL LPS and 5μM Nigericin 100 μL of lysis buffer was added to the cells and incubated at 37 °C for 30 min, debris was pelleted by centrifugation at 10 000 x g for 1 min. Then 50 μL of the reaction mixture was added to the samples and mixed and the samples incubated at room temperature for 20-30 min before adding 50 μL of stop buffer.

### ELISA

IL-1β or TNF in supernatants from THP-1 cells or primary human macrophages were detected using the human IL-1 beta/IL-1F2 DuoSet ELISA (R&D systems DY201-05) or the TNF-alpha DuoSet ELISA (R&D Systems, DY210-05).

### Immunofluorescence staining and imaging

For immunofluorescence experiments, 2 x 10^5^ THP-1 cells/well or 1.0 x 10^6^ PBMCs were seeded in 24-well glass bottom plates (MatTek, P24G-1.5-13-F). Following stimulation, the supernatant was removed, and the cells were fixed either in 1:1 solution of methanol/acetone at-20 °C for a minimum of 20 min followed by rehydration in PBS at RT for 1 h, with 4% paraformaldehyde (PFA) or CytoFix™ (BD BioSciences, 100412100), following the immunostaining protocol previously described. The cells were blocked in 10% A^+^ in PBS for 20 min at RT and immunostained using primary antibodies at 2-5 μg/mL in 1% A^+^ serum in PBS. After washing 3 times for 5 min in PBS the samples were incubated with highly cross-adsorbed fluorescently labeled secondary antibodies at a concentration of 2 μg/mL in 1% A^+^ serum in PBS. The cells were washed 3 times in PBS before incubated for nuclear staining with Hoechst stain (1 μg/ml) for 5 min and washed 2 times in PBS before confocal microscopy. Following fixation all solutions were supplemented with saponin to 0.05 % as previously described (Husebye *et al*., 2010).

Confocal images were captured using a Leica TCS SP8 (Leica Microsystems) equipped with an HC plan-apochromat 63×/1.4 CS2 oil-immersion objective using 488 nm, 561 nm, and 633 nm white laser lines and the 405 nm diode laser for detection. 3-dimensional data was obtained from 12-bit depth imaging data and used to build Z-stacks for individual channels. The white laser intensity was set to 70% as recommended by the manufacturer, and channel fluorescence intensity to 30% or less. Images were also captured on a Nikon Crest v3 Spinning Disk Confocal Microscope equipped using a CFI Plan Apo l D 100X Oil NA 1.45 objective for detection. Imaging was preferentially done between lines and as the same scanning speed and laser intensity for a given set of experiments and with 16-bit depth imaging data. The resulting images enhanced by deconvolution using the NIS-Elements Advanced Research Software Ver.

60.10.01. The Bitplane-IMARIS 8.4.2 software and the 3D-spot function with thresholding were used for quantification giving voxel intensities from imaging raw data. The data was presented as mean voxel intensities.

Using MATLAB, the software could give numerical values that could be analyzed by GraphPad-PRISM 8.2.1. Statistical significance was calculated by One-way or Two-way ANOVA multiple comparison tests, reporting multiplicity adjusted p values (adj. p values) and the results displayed as mean +/-SEM if from three or more independent experiments, or +/-SD if from less than three independent experiments. The statistical tests variants are stated in the figure text.

### Quantitative PCR (qPCR)

6 x 10^5^ THP-1 cells/well or 4.0 x 10^6^ PBMCs were seeded in 6-well plates. Total RNA was isolated from cell lysates in QIAzol (Qiagen, 79306) using chloroform extraction with subsequent purifications on RNeady Mini kit (Qiagen, 74106) following the manufactureŕs protocol. DNA was digested using the DNase set (Qiagen, 79256). RNA concentrations were measured using NanoDrop (Thermo Scientific) and cDNA synthesis was performed using High-Capacity RNA-to-cDNA™ Kit (Thermo Fisher Scientific, 4387406). Quantitative real-time PCR (qPCR) was done using the PerfeCTa qPCR FastMix (Quanta Biosciences, 733-1398) in a 20 μL reaction volume in duplicates and cycled in a StepOnePlusÔ Real-Time PCR cycler (Applied Biosystems, Thermo Fisher Scientific) on technical replicates. The following TaqMan Gene Expression Assays were used; IL-1b (Hs01555410_m1), NLRP3 (Hs00918082_m1), Rab11FIP2 (Hs00208593_m1), Rab11a (Hs00900539_m1), Rab11b (Hs00188448_m1) and TBP (Hs00427620_m1) all from Applied Biosystems, Thermo Fisher Scientific).

### DNA transfection and co-immunoprecipitation

List of plasmids used: hsNLRP3 in pEGFP-C2 was a gift from Christian Stehlik (Addgene plasmid # 73955; http://n2t.net/addgene:73955; RRID:Addgene_73955) and hsASC-HA in pCI was a gift from Kate Fitzgerald (Addgene plasmid # 41553; http://n2t.net/addgene:41553; RRID:Addgene_41553), hsFIP2 in pLVX (Skjesol *et al*., 2019) and hsRab11 in pECFP-C1 (Husebye *et al*., 2010). HEK293T cells were seeded at 250 000 cells/well in 6-well plates and transfected the following day using Genejuice transfection reagent (GeneJuice® Transfection Reagent, 70967-5). 48 h after transfection, cells were in PBS and harvested in ice-cold lysis buffer (150 mM NaCl, 50 mM Tris-HCl pH 8.0, 1 mM EDTA, 1% NP-40) with added 50 mM NaF, 2 mM Na_3_VO_3_ (Sigma-Aldrich) containing PhosSTOP phosphatase inhibitor cocktail (Roche, 4906845001) and cOmplete mini protease inhibitor cocktail (Roche, 11836153001).

Lysates were cleared by centrifugation at 21000 x g at 4 °C for 15 min, and co-IPs were performed using 50 μL anti-Flag (M2) affinity agarose (Sigma-Aldrich, A2220) on rotation at 4 °C for 3 h or ON. Samples were washed four times and eluted at 80 °C for 7 min in 1x NuPAGE LDS sample buffer (Thermo Fisher Scientific, NP0007), and agarose beads were subsequently removed by high-speed centrifugation. DTT was added to a concentration of 25 mM and samples were heated again (80 °C for 7 min) before being subjected to SDS-PAGE and Western blotting.

### SDS-PAGE and Western Blotting

Protein was precipitated by isopropanol precipitation from the organic phase of Qiazol lysates, after RNA samples had been isolated, and according to manufacturer’s instructions (Thermo Fisher Scientific). The pellet was air dried and dissolved in 4 M urea in 2 % SDS before heated at 95 °C dissolved in 1x LDS supplemented with 25 mM DTT for 5 min. Samples were loaded and run on 4-12 % Bis-Tris NuPage gels and blotted on nitrocellulose membranes using the iBlot2 system (Thermo Scientific, IB21001). Membranes were blocked in 5% BSA for 1 h at RT before incubation with primary antibodies up to 72 h. Membranes were washed in 3 times in TBS-T (Tris Buffered Saline with 0.1% Tween-X100) before incubated with HRP-conjugated immunoglobulins for 1 h at RT. The following HRP-conjugated immunoglobulins from Agilent Dako Products were used: Swine anti-rabbit immunoglobulins/HRP (P039901-2), Goat anti-mouse immunoglobulins/HRP (P044701-2) and rabbit anti-goat immunoglobulins/HRP (P044901-2). Following 3 washes in TBS-T the blots were developed using SuperSignal West Femto Substrate (Thermo Scientific, 34095). Images were captured with the Li-COR Odyssey system and analyzed by the Odyssey Image Analysis software Ver 5.2.

## Acknowledgments

The confocal imaging was performed at the Cellular and Molecular Imaging Core Facility (CMIC), Norwegian University of Science and Technology (NTNU). CMIC is funded by the Faculty of Medicine and Health Sciences at NTNU and Central Norway Regional Health Authority. The Research Council of Norway through its Centers of Excellence funding scheme, awarded the grant #223255/F50 to TE, that financed: CG, MY, AS, CC, TE and HH. The Research Council of Norway, FRIMEDBIO program, awarded the grant # 275876 to TE, that financed: CG, MY, AS, CC, ZI, CRD, TE and HH. The Research Council of Norway, FRIMEDBIO program, awarded the grant #100412100 to TE, that financed: SK, LR, TE and HH.

## Author Contributions

Caroline Gravastrand: Methodology; data curation; formal analysis; validation; investigation, writing review and editing.

Mariia Yurchenko: data curation, formal analysis, validation, investigation, review and editing.

Stine Kristensen: Methodology; data curation; formal analysis; validation, investigation, editing.

Astrid Skjesol: Methodology; data curation; validation; investigation

Carmen Chen, Zunaira Iqbal, Karoline Ruud Dahlen, Unni Nonstad and Liv Ryan: Methodology

Terje Espevik: Conceptualization; investigation, data curation, funding acquisition; writing – review and editing.

Harald Husebye: Conceptualization; supervision; project administration; methodology, data curation; formal analysis; validation; investigation; visualization, writing – review and editing.

## Expanded View Figure legends

**Figure EV1.**
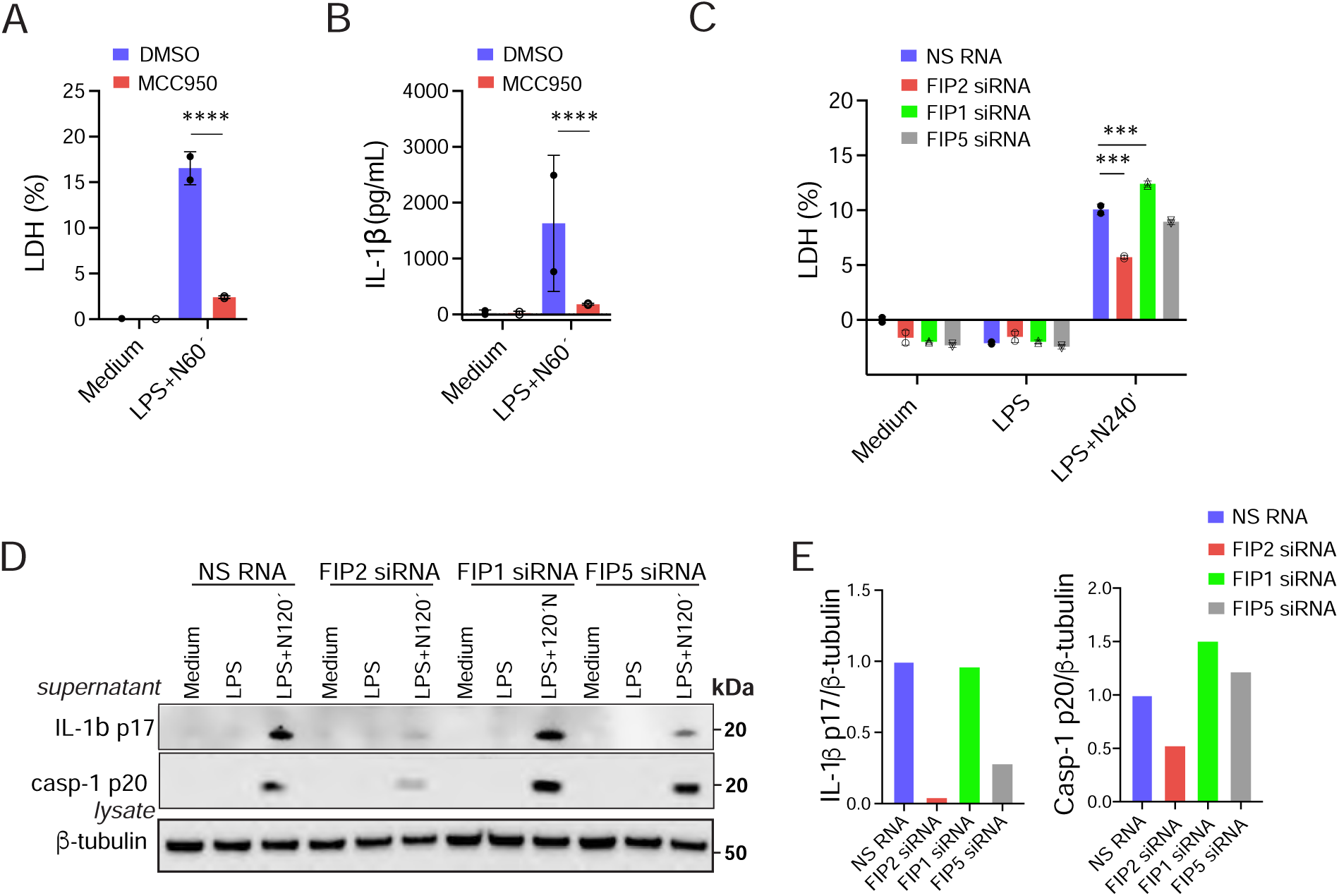
FIP2 is the driver of caspase-1 mediated IL-1β cleavage of the class I FIPs (**A**) Quantification of LDH release in THP-1 cells treated with DMSO or NLRP3 inhibitor MCC950 before LPS priming and nigericin treatment, n = 2 independent experiments. (**B**) IL-1β ELISA from the cell supernatants in A. (**C**) Quantification of LDH-release in primary human macrophages treated with NS RNA, FIP1 siRNA, FIP2 siRNA or FIP5 siRNA. (**D**) Immunoblot of pro-IL-1β, IL-1β p17, pro-caspase-1, and caspase-1 p20 in lysates from the FIP silenced Mφ‘s in C. (**E**) Quantification of IL-1β p17 and caspase-1 p20 levels normalized towards β-tubulin in the immunoblots from D. The cells were treated with respective siRNÁs before primed with 100 ng/mL LPS for 2 h and treated with 5 μM Nigericin as indicated, one experiment, n = 1. In (A-C) data are presented a as mean +/-SD. ** p=0.0047, *** P=0.0002 (Two-way ANOVA Tukey’s multiple comparisons test with adj. p values).

**Figure EV3.**
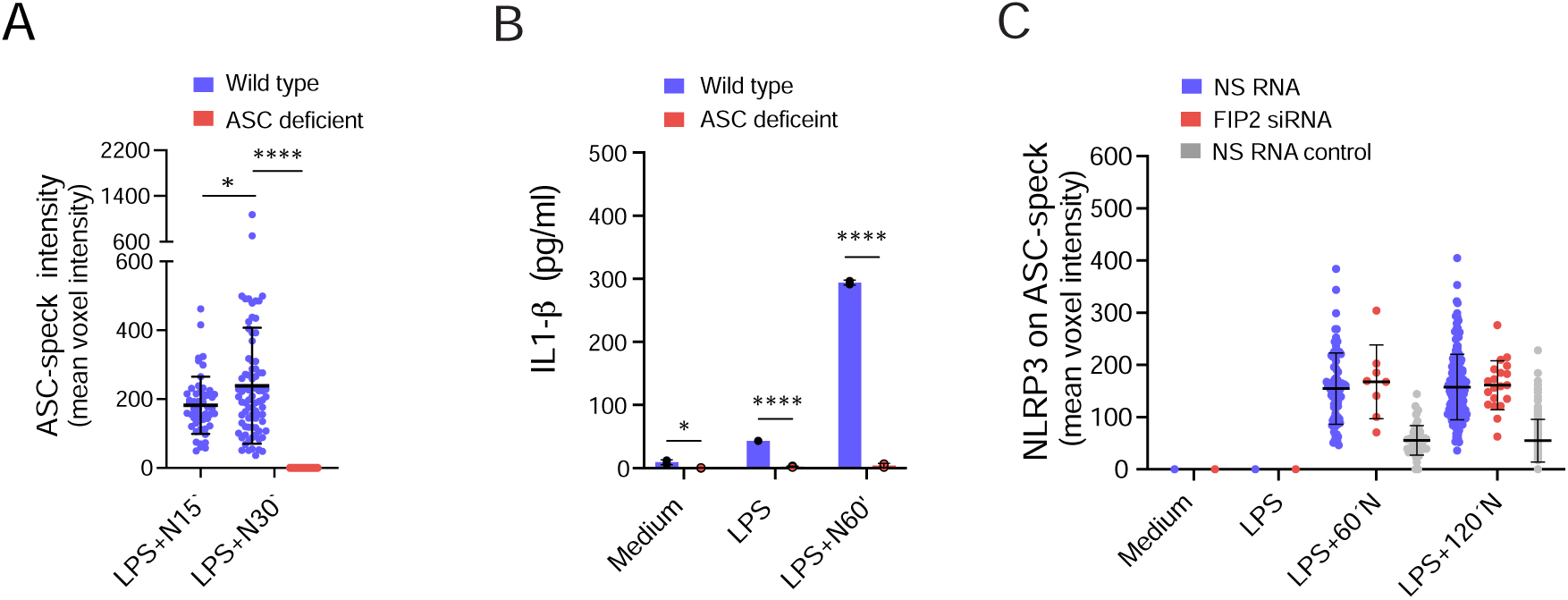
Characterization of ASC-deficient cells and NLRP3 intensity on the ASC-speck (**A**) Quantification of ASC-speck intensity in wild type and ASC-deficient THP-1 cells. 266-483 cells were monitored per condition. (**B**) Quantification of IL-1β release in wild type and ASC-deficient THP-1 cells by ELISA. (**C**) Quantification of NLRP3 on ASC-speck in NS RNA or FIP2 siRNA treated human macrophages from the donor presented in Figure 3F. ASC-specks were identified by using the spot detection mode of the IMARIS 8.2 imaging software on 3-D confocal imaging raw data. In (A-C) data are presented a as mean +/-SD. * p=0.0155-0.0450, *** p=0.0003 and **** p < 0.0001 (One-way ANOVA with adj. p values). N = nigericin. Mean +/-SD.

**Figure EV5.**
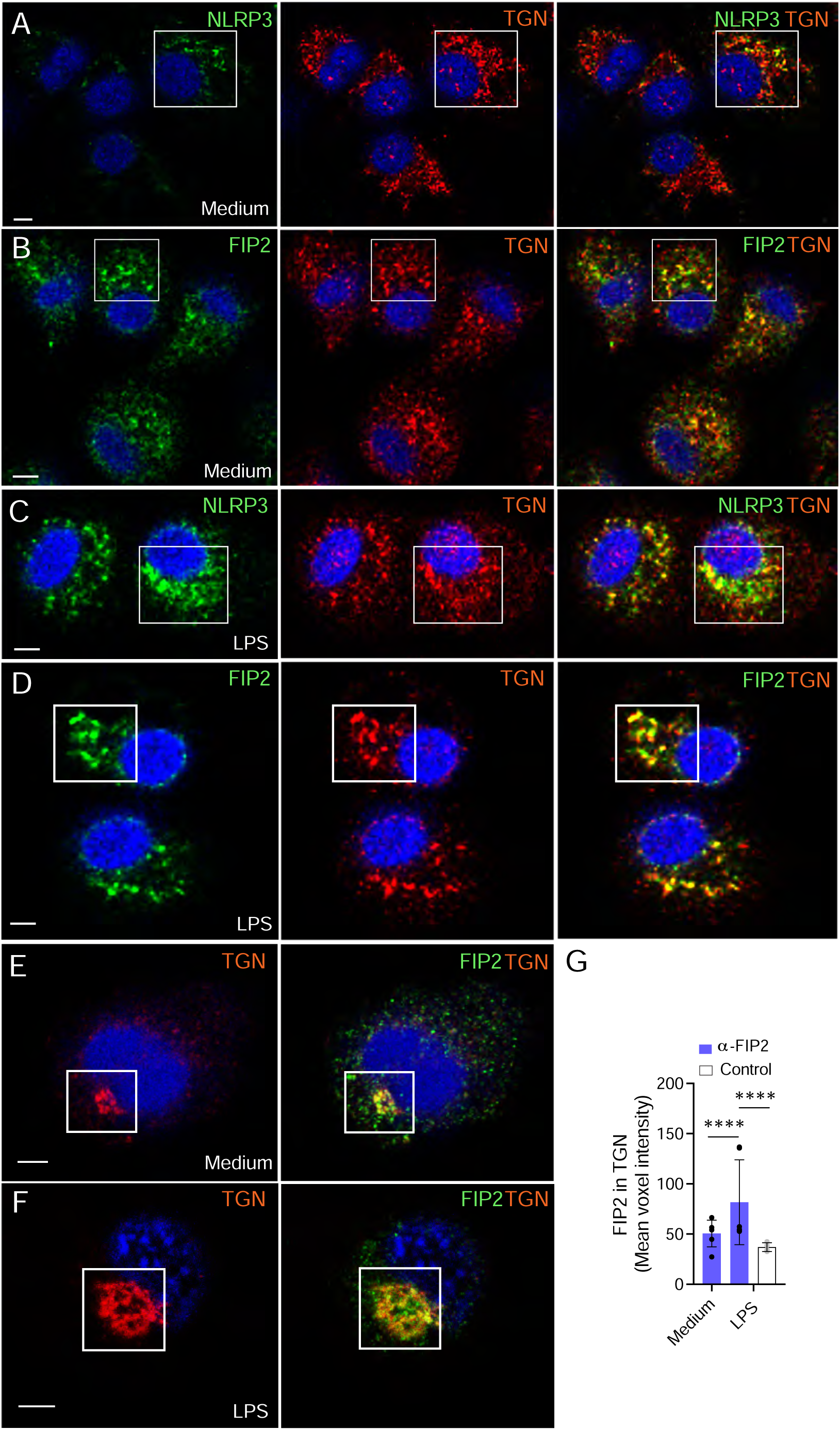
FIP2 and NLRP3 both localize to the TGN in primary human macrophages (**A**) Confocal image showing NLRP3 (green) and TGN46 (red) in the TGN of unstimulated macrophages. (**B**) Confocal image showing FIP2 (green) and TGN46 (red) in the TGN of unstimulated macrophages. (**C**) Confocal image showing NLRP3 (green) in the TGN46 (red) of LPS primed macrophages. (**D**) Confocal image showing FIP2 (green) in the TGN46 (red) of LPS primed macrophages. (**E**) Confocal image showing FIP2 (green) and the FIP2/TGN46 overlay in the TGN of unstimulated THP-1 cells. (**F**) Confocal image showing FIP2 (green) and FIP2/TGN46 overlay in the TGN of LPS primed THP-1 cells. **(G**) Quantification of FIP2 in the TGN-ring structure of unstimulated and LPS primed cells. The TGN46 positive peri-nuclear ring and dTGN structures were identified using the IMARIS 8.2 imaging software on 3-D confocal imaging raw data. The cells were left unstimulated of LPS primed for 2 h before fixation and co-immunostained with FIP2 and TGN46 antibodies. The data (G) is presented as mean +/-SD. ** (p = 0.0072) and **** (p<0.0001) (Two-way ANOVA multiple comparisons test with adj. p values). Arrows - point at TGN46 positive dTGN.Scale bar = 5 μm.

**Figure EV6.**
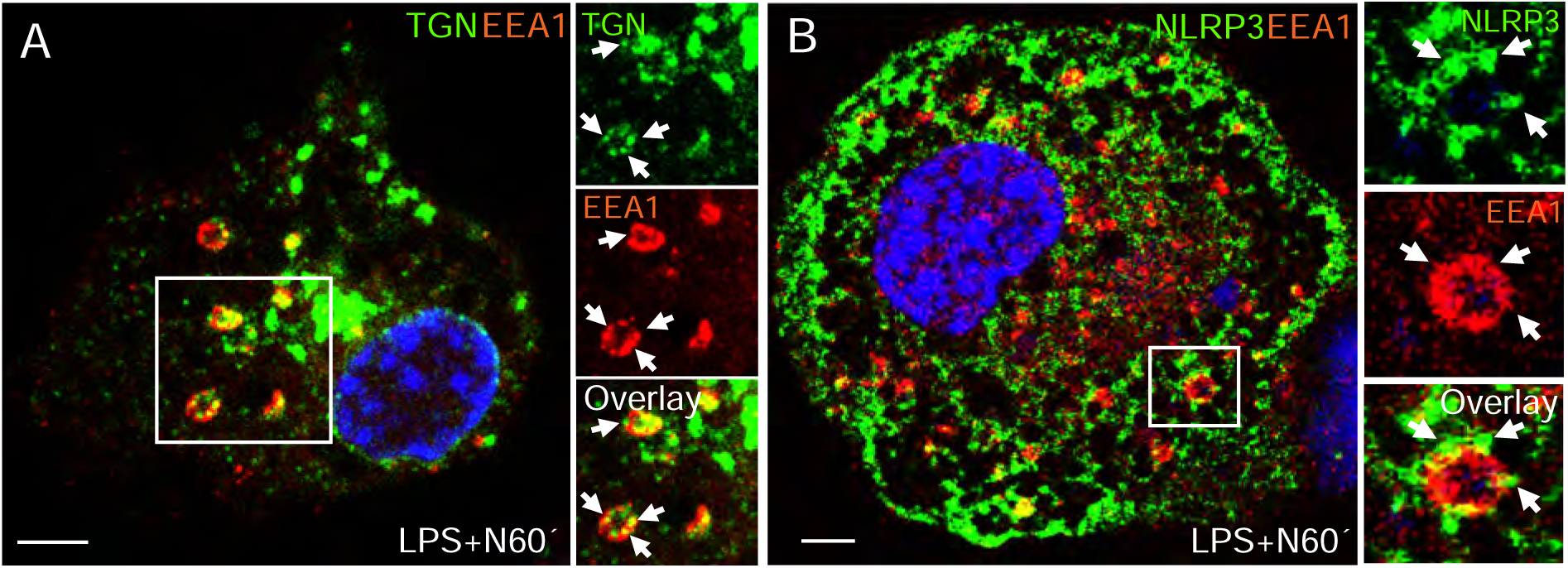
TGN46 and NLRP3 are locate to microdomains on EEA1 positive endosomes (**A**) Confocal images showing dTGN structures (green) positive for EEA1 (red). (**B**) Confocal images showing NLRP3 (green) on a EEA1 (red) positive endosome. The cells were primed with 100 ng/mL LPS for 2 h and treated with 5 μM nigericin for 1 h. Scale bar = 5 μm. Arrows points at TGN46 or NLRP3 microdomains on enlarged EEA1 positive endosomes.

**Figure EV7.**
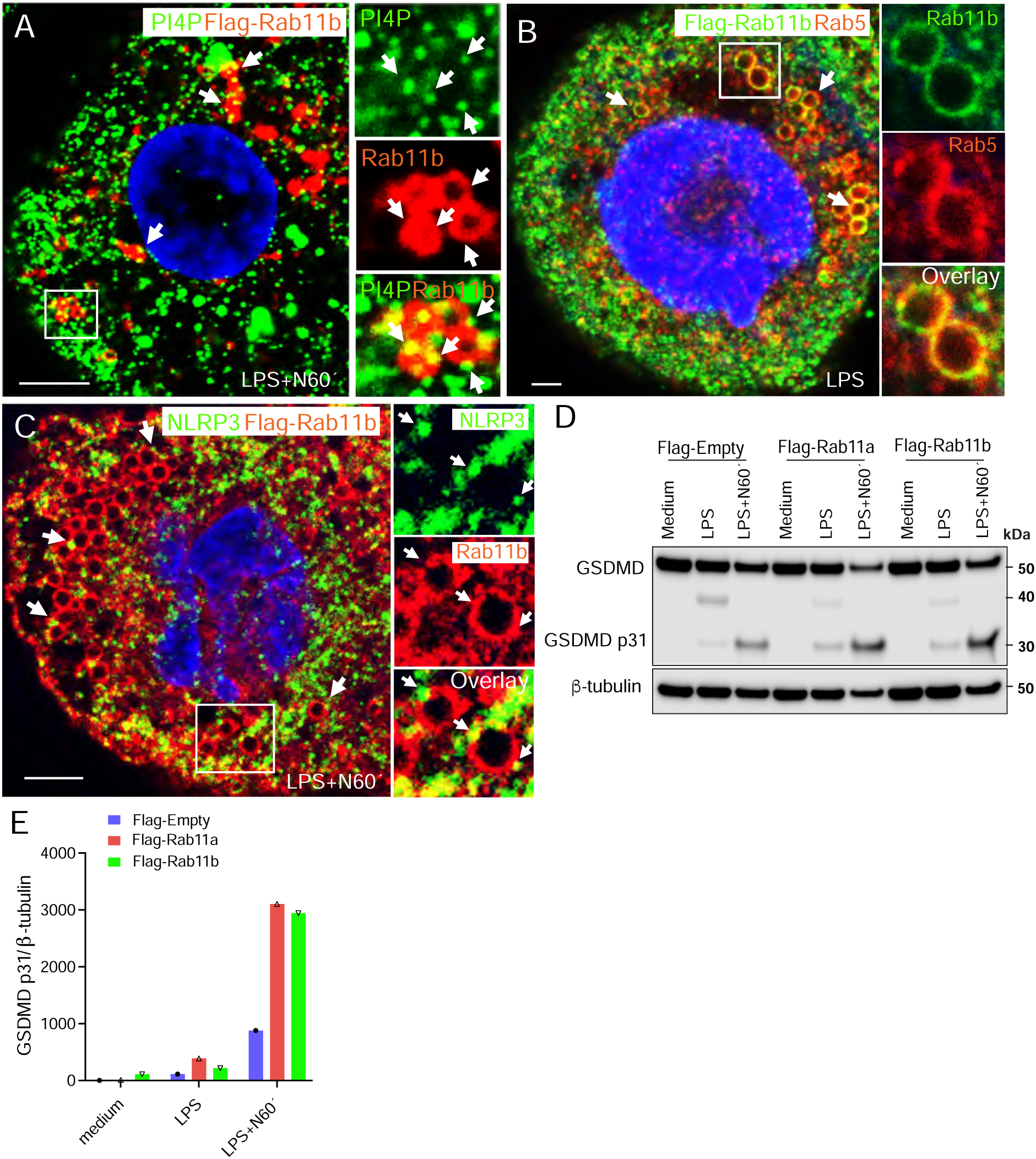
PI4P and NLRP3 are locate to microdomains on Rab11b positive early endosomes. (**A**) Confocal image showing Rab5 (red) on Rab11b positive endosomes (green) in Flag-Rab11b expressing THP-1 cells. (**B**) Confocal image showing NLRP3 (green) positive microdomains on Flag-Rab11b endosomes (red). (**C**) Confocal image showing PI4P (green) positive microdomains on Flag-Rab11b positive endosomes. (**D**) Immunoblot total GSDMD, caspase-1 cleaved p31 GSDMD fragment and β-tubulin in Flag-Empty, Flag-Rab11a and Flag-Rab11b co-expressing THP-1 cells. (**E**) Quantification of the GSDMD p31 fragment in the cells from (D). The cells were primed with 100 ng/mL LPS for 2 h and treated with 5 μM nigericin for 1 h. Scale bar = 5 μm.

